# A versatile miniature two-photon microscope enabling multicolor deep-brain imaging

**DOI:** 10.1101/2025.02.13.635171

**Authors:** Runlong Wu, Chunzhu Zhao, Shan Qiu, Yufei Zhu, Lifeng Zhang, Qiang Fu, Yanhui Hu, Dakun Wu, Fei Yu, Fangxu Zhou, Haipeng Huang, Yunfeng Zhang, Xianhua Wang, Aiming Wang, Heping Cheng

**Affiliations:** National Biomedical Imaging Center, State Key Laboratory of Membrane Biology, Institute of Molecular Medicine, Peking-Tsinghua Center for Life Sciences, College of Future Technology, Peking University, Beijing, China; Beijing Key Laboratory for Optoelectronics Measurement Technology, Beijing Information Science and Technology University, Beijing, China; Beijing Laboratory of Biomedical Imaging, Beijing Municipal Education Commission, Beijing, China; Academy for Advanced Interdisciplinary Studies, Peking University, Beijing, China; Research Unit of Mitochondria in Brain Diseases, Chinese Academy of Medical Sciences, PKU-Nanjing Institute of Translational Medicine, Nanjing Raygen Health, Nanjing, China; Beijing Transcend Vivoscope Biotech Co., Ltd., Beijing, China; Hangzhou Institute for Advanced Study, University of Chinese Academy of Sciences, Hangzhou, China; Key Laboratory of Materials for High Power Laser, Shanghai Institute of Optics and Fine Mechanics, Chinese Academy of Sciences, Shanghai, China; School of Electronics, Peking University, Beijing, China; State Key Laboratory of Advanced Optical Communication System and Networks, Peking University, Beijing, China

## Abstract

Here we present the FHIRM-TPM 3.0, a 2.6 g miniature two-photon microscope capable of multicolor deep-brain imaging in freely behaving mice. The system was integrated with a broadband anti-resonant hollow-core fiber featuring low transmission loss, minimal dispersion from 700-1060 nm, and high tolerance of laser power. By correcting chromatic and spherical aberrations and optimizing the fluorescence collection aperture, we achieved cortical neuronal imaging at depths exceeding 820 μm and, using a GRIN lens, hippocampal Ca^2+^ imaging at single dendritic spine resolution. Moreover, we engineered three interchangeable parfocal objectives, allowing for a tenfold scalable field-of-view up to 1×0.8 mm², with lateral resolutions ranging from 0.68 to 1.46 μm. By multicolor imaging at excitation wavelengths of 780 nm, 920 nm and 1030 nm, we investigated mitochondrial and cytosolic Ca^2+^ activities relative to the deposition of amyloid plaques in the cortex of awake APP/PS1 transgenic mice. Thus, the FHIRM-TPM 3.0 provides a versatile imaging system suitable for diverse brain imaging scenarios.

---

The development of miniature microscopes has been instrumental in deciphering the neural basis of behaviors in naturally behaving animals. While single-photon fluorescence microscopes [1–4] have been valuable tools, multiphoton imaging affords unique advantages, such as inherent optical sectioning and deep tissue penetration, making it suitable for volumetric or multi-plane recording at the levels of synapses, neurons and local circuities. Building upon long-standing efforts [5–8], the recent advent of miniaturized two-photon microscopes (m2PMs) [9–12] and miniature three-photon microscopes (m3PMs) [13, 14] has empowered research in experimental paradigms that were previously impossible with physically constrained animals. For instance, studies in freely behaving animals have uncovered a dynamic disinhibitory microcircuit involved in social competition [15], sparse GABAergic neural ensembles encoding social novelty [16], spatial tuning of neurons during unrestrained behaviors [11, 17], and dynamic surveillance of microglia during wake-sleep cycles [18].

Despite these advancements, current m2PM technologies still face several unmet challenges, particularly in multicolor fluorescence imaging, achieving greater imaging depth, and attaining a larger, scalable field of view (FOV). In prevalent m2PM and m3PM systems, the delivery of femtosecond laser pulses relies on photonic bandgap hollow-core fibers (PBG-HCF), each transmitting laser only at specific wavelengths. Consequently, dual-color imaging in recent reports has exploited a single laser to excite two different indicators, e.g., 920 nm for GCaMP6 and tdTomato [11], albeit with compromised efficiency [19–21]. In a multicolor two-photon endomicroscopy study, Guan et al. employed a double-clad fiber to deliver 830 nm and 1050 nm wavelengths, while a third virtual wavelength at 927 nm was generated by synchronizing two coherent pulses, enabling imaging of four fluorophores (CFP, GFP, YFP and RFP) simultaneously [22]. However, the system’s FOV of 120 μm and frame rate of 1.5 Hz need to be improved for functional imaging of large cohorts of neurons.

In state-of-the-art benchtop systems, two-photon microscopy can achieve cortical imaging depths exceeding 800 μm [23], while three-photon microscopy can reach depths up to 1.4 mm [24] for neuronal Ca^2+^ activity imaging. Remarkably, a recently developed m3PM system [13] reached a depth of 1.2 mm, enabling hippocampal CA1 Ca^2+^ imaging in freely moving mice. This result is inspiring because it demonstrates that a miniature multiphoton system can approach the imaging depths of its benchtop counterparts. Since previously reported m2PM imaging depths were mostly around 250 μm without a GRIN lens [9–12, 22, 25, 26], further development of m2PM is warranted to unleash its full potential for deep brain imaging in freely behaving animals.

As for the FOV of m2PM, it has been steadily increased from 130×130 μm^2^ in FHIRM-TPM [9] to 420×420 μm^2^ in FHIRM-TPM 2.0 [10], and more recently, even to 1×0.788 mm^2^ [26]. Furthermore, by analogy to benchtop systems, the advantage of an optically scalable FOV would help balance resolution requirements with the size of neuronal cohorts being surveyed, enabling m2PM to investigate the same area at different magnifications. Clearly, overcoming the aforementioned limitations collectively would significantly enhance the applicability of m2PM technology.

Here we have developed the FHIRM–TPM 3.0, a versatile m2PM system incorporating multiple desirable features on top of previous developments. By designing and fabricating a broadband anti-resonant hollow-core fiber (AR-HCF) and systematically optimizing the entire optical design, we achieved multicolor imaging at three excitation wavelengths (780 nm, 920 nm, and 1030 nm), and extended the imaging depth to over 820 μm in the cortex. Furthermore, by switching among three interchangeable parfocal objectives of different magnifications, our microscope offers a tenfold change in FOV, providing flexibility to accommodate various experimental requirements.

## Results

### The design of the FHIRM-TPM 3.0

The overall configuration of the FHIRM-TPM 3.0 system was consistent with that reported in our previously developed microscopes [9, 10], incorporating important improvements and modifications (**Fig. 1a, b, Extended Data Fig. 1a**). In our setup, we employed three femtosecond fiber lasers with central wavelengths of 780 nm, 920 nm, and 1030 nm to achieve simultaneous three-color excitation (**Extended Data Fig. 1a**). To minimize the size and weight of the headpiece and reduce co-location errors caused by the convergence of multiple fiber output paths, we aimed to deliver multiple laser wavelengths through a single optical fiber. The PBG-HCF used in previous m2PM systems [9–11] had a relatively narrow bandwidth (<100 nm) and high dispersion (∼75 fs/nm/m at 920 nm) [9, 10]. In this regard, recently developed AR-HCFs [13, 27, 28], despite their wider transmission bandwidth (370-700 nm or 1100-1700 nm), cannot be directly used for two-photon imaging with excitation bandwidth of 700-1100 nm. To address these limitations, we developed a new broadband AR-HCF. By designing a seven-tube cladding with extremely thin silica walls, this fiber achieved low transmission loss (< 0.2 dB/m) and reduced dispersion, estimated to be 1.53-2.70 fs/nm/m, over the wavelength band range from 700 nm to 1060 nm (**Fig. 1c, Supplementary table 1**). The bending loss was negligible when the bend radius exceeded 4 cm (**Extended Data Fig. 2a, b**). The Gaussian beam profiles exhibited the quasi-single-mode guidance of this fiber (**Extended Data Fig. 2c**). The relatively large fiber core diameter of 28 μm also ensured a high damage threshold. The test data with the 920 nm laser at power greater than 1.5 W demonstrated that the output laser power remained stable for more than 24 hours, with a peak-to-peak power fluctuation of approximately 0.72% (**Extended Data Fig. 2d**). The fiber maintained high fidelity to the pulse waveform. Pulse durations measured at the exit of a 2-meter AR-HCF were 167±2 fs, 139±2 fs, and 136±3 fs for 780 nm, 920 nm, and 1030 nm lasers, respectively, and were close to the respective input pulse widths, with minimal distortion observed when the power was adjusted from 50 to 700 mW (**Fig. 1d**).

**Figure 1.**
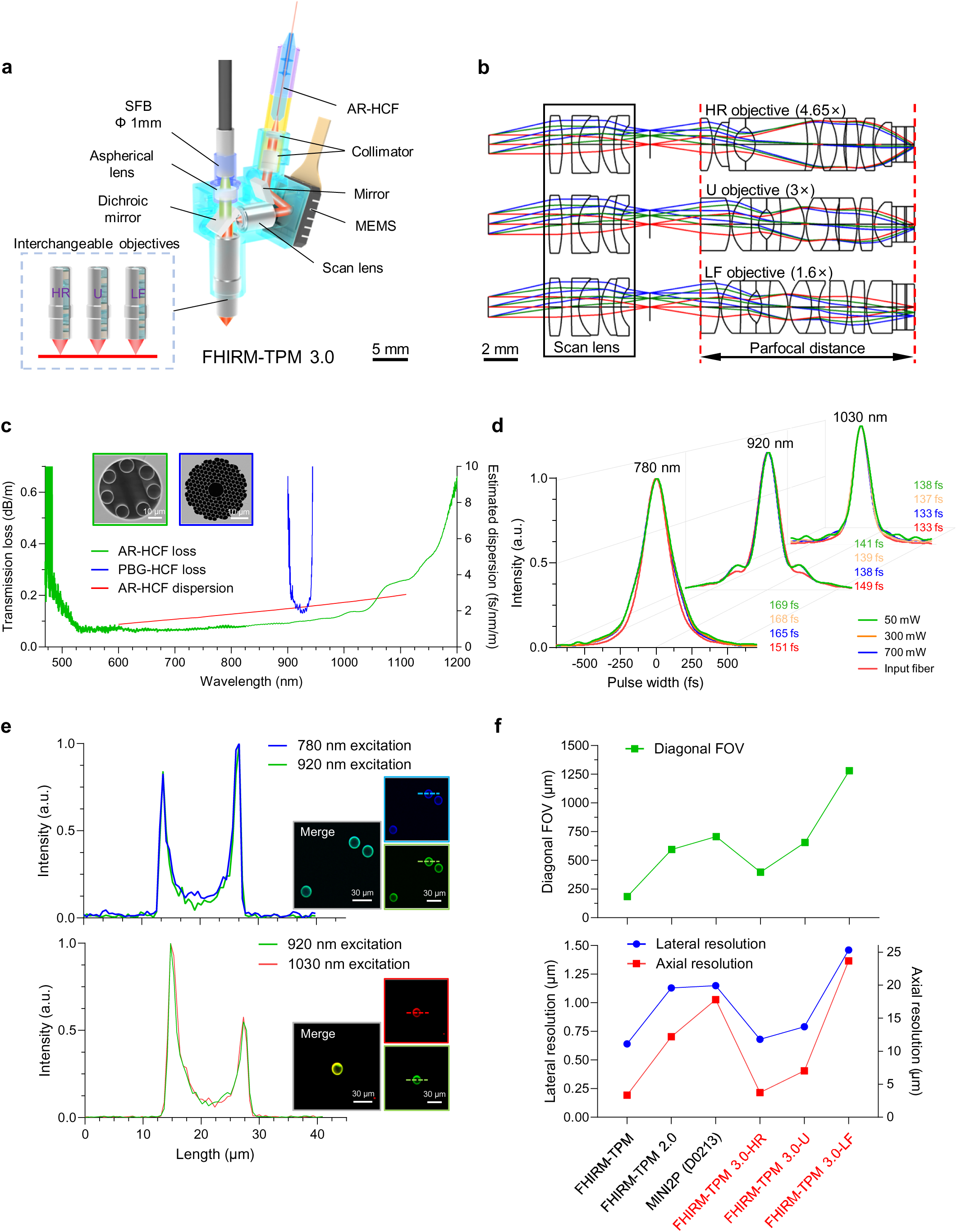
Design and test of FHIRM-TPM 3.0. **(a)** Schematic of the headpiece. AR-HCF, anti-resonant hollow-core fiber, for 700 nm-1100 nm waveband transmission; MEMS, micro-electro-mechanical systems; SFB, supple fiber bundle. Three interchangeable objectives of different magnification: HR, high resolution; U, universal; LF, large field of view (FOV). **(b)** Optical design. A scan lens with a large scan angle is shared by the HR, U, and LF objectives that have the same parfocal distance. **(c)** Transmission loss and estimated dispersion of the AR-HCF and a customized photonic bandgap hollow core fiber (PBG-HCF). Inset, electron micrographs of fiber cross-sections. **(d)** Autocorrelation profiles, showing the pulse widths of 780 nm, 920 nm and 1030 nm femtosecond lasers before and after transmitting through a 2-m AR-HCF at different power levels. A.u., arbitrary units. **(e)** Chromatic aberration tests of the FHIRM-TPM 3.0-U headpiece. Top, the plot shows intensity profiles along the cross lines in the inset on the right. Inset, images of 15 μm beads at 780 nm (upper right), 920 nm excitation (lower right), and the merged image (left). Bottom, the same as above, except for excitation wavelengths of 920 nm and 1030 nm. **(f)** Comparison of diagonal FOVs (top), and lateral and axial resolutions (bottom) of different m2PM systems [9–11].

To accommodate the requirements for both multicolor imaging and large, multiscale FOVs, we have engineered three sets of objectives of different magnifications. These included a high-resolution objective (HR, 4.65×, excitation NA: 0.7), a universal utility objective (U, 3×, excitation NA: 0.5) and a large FOV objective (LF, 1.6×, excitation NA: 0.26) **(Fig. 1b).** All three objectives shared the same large-angle scan lens, parfocal distance, outer diameter and headpiece-mounting procedures, making them fully interchangeable and allowing for the investigation of the same region of interest at multiscale FOVs and resolutions.

Importantly, the entire optical system, including the collimator, scan lens, and all objectives, was systematically designed to achieve chromatic aberration correction across wavebands from 760 nm to 1100 nm (**Extended Data Fig. 3a-f, Supplementary Note 1**). Simulation results showed that residual lateral chromatic aberrations across this range were less than 0.5 μm for FHIRM-TPM 3.0-HR/U/LF (**Extended Data Fig. 3b**). Residual axial chromatic aberrations were approximately 1 μm for FHIRM-TPM 3.0-HR/U and around 3 μm for FHIRM-TPM 3.0-LF (**Extended Data Fig. 3c**), much smaller than the respective diffraction limits in the axial direction. Testing with 15 μm fluorescent beads using FHIRM-TPM 3.0-U under 780 nm, 920 nm, and 1030 nm excitation confirmed a good achromatism in our system (**Fig. 1e**). For deep brain imaging, we optimized the system design to reduce spherical aberrations, thereby increasing two-photon excitation efficiency (**Extended Data Fig. 3d**). Additionally, we expanded the collection NAs for the three objectives to more efficiently collect scattered fluorescence from deep brain tissues (**Extended Data Fig. 3e, f, Supplementary Note 1**).

Using 920 nm for excitation, FHIRM-TPM 3.0-HR achieved spatial resolutions of 0.68 μm laterally and 3.73 μm axially over a FOV of 300×260 μm^2^ (**Fig. 1f, Supplementary Table 2**). With a FOV of 500×425 μm^2^, FHIRM-TPM 3.0-U achieved a lateral resolution of 0.79 μm and an axial resolution of 7.04 μm, a significant improvement compared to FHIRM-TPM 2.0 [10] (**Fig. 1f, Extended Data Fig. 4a, b, Supplementary Table 2**). By integrating the strategies recently developed to enlarge the FOV [25], the FHIRM-TPM 3.0-LF achieved a millimeter-scale FOV (1×0.8 mm^2^) while maintaining resolutions of 1.46 μm laterally and 23.68 μm axially (**Fig. 1f, Supplementary Table 2**).

### Expanding the scope of in vivo imaging with FHIRM-TPM 3.0

Next, we sought to demonstrate FHIRM-TPM 3.0’s capabilities for in vivo functional and structural brain imaging in head-fixed and freely moving animals (**Supplementary Tables 3** and **4**). To evaluate aberrations in dual-color Ca²⁺ imaging, neurons co-expressing GCaMP6f and jRGECO1a in M1 were alternately illuminated at 920 nm and 1030 nm (5 Hz), and we found that the Ca^2+^ signals from dual-color excitation overlapped in space and time with little distortion (**Extended Data Fig. 5**). In mice with GCaMP6s-labeled neurons and mCherry-labeled astrocytes in the M1, FHIRM-TPM 3.0-U was applied for dual-color imaging with simultaneous 920 nm and 1030 nm excitation (**Extended Data Fig. 6a, b, Supplementary Video 1**). The robustness of the imaging system was demonstrated by its ability to continuously track neuronal Ca^2+^ transients and the morphology of astrocytes, even during sudden and vigorous movements of the animal in response to an electric foot shock. Furthermore, dual-color Ca^2+^ signals from GCaMP6s- and jRGECO1a-labelled M1 neurons were recorded during a 30-minute exploration session in an 80-cm square open-field box (**Extended Data Fig. 6c-e**).

To assess its deep-brain imaging capability, we used FHIRM-TPM 3.0-U to acquire 854 μm z-stacks in the M1 cortex of awake, head-fixed Thy1-YFPH transgenic mice (**Fig. 2a, Supplementary Video 2**). For dual-wavelength excitation of an indicator at a depth of 850 μm, the laser power under the objective was 110 mW for 920 nm and 67 mW for 1030 nm (**Fig. 2b, c**). Additionally, a Ca^2+^ imaging stack in awake, head-fixed mice expressing GCaMP6s attained a depth of 822 μm (**Extended data Fig. 7a, b, Supplementary Video 3**) and time-lapse single-plane imaging of Ca^2+^ transients was performed at 500 μm depth with a high signal-to-noise ratio (SNR) at 10 Hz under freely moving conditions (**Extended data Fig. 7c, d, Supplementary Video 4**). In addition, we were able to resolve Ca^2+^ activities in individual dendritic spines of hippocampus CA1 neurons using FHIRM-TPM 3.0-U through a GRIN lens (1-mm diameter, 4-mm length), despite the aberrations it introduced (**Extended Data Fig. 8a-c, Supplementary Note 2, Supplementary Video 5**).

**Figure 2.**
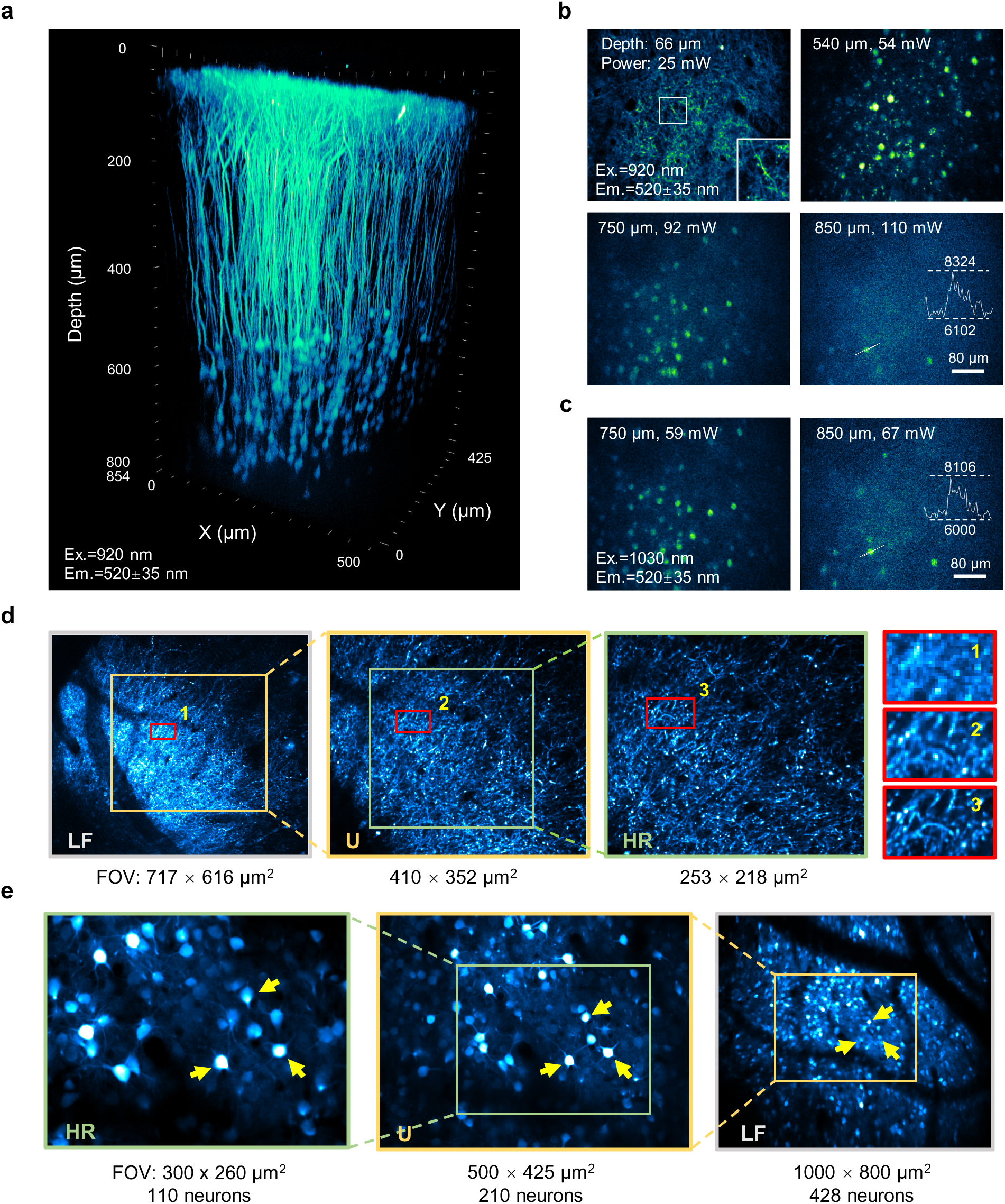
FHIRM-TPM 3.0 achieves deep-cortex and scalable FOV imaging in vivo. **(a)** Reconstruction of an 854 μm z-stack in the M1 of a Thy1-YFPH transgenic mouse using 920 nm excitation with FHIRM-TPM 3.0-U. Ex., excitation; Em., emission. **(b, c)** Representative 7-frame-averaged x-y images at designated depths using 920 nm **(b)** and 1030 **(c)** nm excitation, respectively. At the 850-μm depth, the images were smoothed using a Gaussian kernel (σ = 1 pixel), with insets showing the intensity profiles along the cross lines. The power values refer to the average laser power after the miniature objective. **(d, e)** Imaging the same area with different objectives. See Methods for details of changing the objectives. In awake mice, GCaMP6f-labeled mitochondria in the M1 region at a depth of 60 μm **(d)** and GCaMP6s-labeled neurons in the mPFC region at a depth of 200 μm **(e)** were imaged using HR, U, and LF objectives, respectively, with image resolution **(d)** and FOV **(e)** progressively increasing from left to right. Rightmost inset in **d** shows the same mitochondrial structures (marked with numerals in the images) under the three objectives. Yellow arrows in **e** show the same three neurons at different optical magnifications, with FOV varying from 300×260 μm^2^ to 1000×800 μm^2^. Data in **a, b** and **c** are representative of three experiments from two mice. Data in **d** and **e** are representative of three experiments from three mice.

To demonstrate the scalable imaging capability of FHIRM-TPM 3.0, we achieved a tenfold expansion in FOV balanced with sufficient resolutions by switching from the HR to the LF objective, with the same region-of-interest and focal plane readily relocated (**Fig. 2d, e**). For visualizing fine structures, FHIRM-TPM 3.0-HR exhibited a high SNR when imaging GCaMP6f-labled mitochondria in the dendritic arbors of M1 neurons (**Fig. 2d**). We reconstructed sparsely labeled mitochondrial networks in a neuron (**Extended Data Fig. 9a, Supplementary Video 6**). Additionally, FHIRM TPM 3.0-HR was applied to track the movement of individual mitochondria and mitochondrial fission-fusion events in vivo (**Extended Data Fig. 9b, Supplementary Video 7**). Utilizing FHIRM-TPM 3.0-LF, we realized large-FOV deep-cortex imaging of 1000×800×702 μm^3^ of the medial prefrontal cortex (**Extended Data Fig. 10a, Supplementary Video 8**). We also acquired Ca^2+^ signals from more than 400 neurons on single selected focal planes in freely moving animals (**Extended Data Fig. 10b, c, Supplementary Video 9**), demonstrating feasibility of simultaneous Ca²⁺ imaging in hundreds of neurons.

### Multicolor imaging of mitochondrial and cytosolic Ca^2+^ and amyloid plaques in the cortex of APP/PS1 transgenic mice

Previous studies have shown that clusters of hyperactive neurons and neurite Ca^2+^ overload occur near amyloid plaques in the cortex of APP23xPS45 or APPswe/PS1-ΔE9 double transgenic Alzheimer’s disease (AD) mice, as early as 6 months old [29, 30]. Because mitochondrial Ca^2+^ ([Ca^2+^]_mito_) is crucial for bioenergetics and cell signaling and is regulated by cytosolic Ca^2+^ ([Ca^2+^]_cyto_) [31, 32], our aim is to determine whether and how cortical [Ca^2+^]_mito_ dysregulates during AD progression and, if so, its relation with [Ca^2+^]_cyto_ change and the plaque disposition.

In excitatory neurons of the M1 cortex of APP/PS1 mice and age-matched wild-type (WT) controls, we expressed two genetically encoded Ca^2+^ indicators, jRGECO1a and GCaMP6f, targeting the cytosolic and mitochondrial compartments and subject to two-photon excitation at 1030 nm and 920 nm, respectively. In vivo visualization of amyloid plaques was achieved using methoxy-X04 [33] with two-photon excitation at 780 nm (**Fig. 3a**). Leveraging the multicolor imaging ability provided by the FHIRM-TPM 3.0-U, we could reconstruct the 3D architecture of mitochondria and neurons relative to sites of amyloid plaques (**Fig. 3b, Supplementary Video 10**). We then recorded [Ca^2+^]_mito_ and [Ca^2+^]_cyto_ transients at selected focal planes within Layer 1 or Layer 2/3 by time-lapse imaging. **Fig. 3c** shows representative traces of simultaneously acquired [Ca^2+^]_mito_ and [Ca^2+^]_cyto_ transients in neurites and somata. Consistent with previous work [34], the majority of [Ca^2+^]_cyto_ transients occurred without [Ca^2+^]_mito_ increases while [Ca^2+^]_mito_ transients were always triggered by an ongoing [Ca^2+^]_cyto_, indicating a loose [Ca^2+^]_mito_ and [Ca^2+^]_cyto_ coupling. We found that elevated [Ca^2+^]_mito_ activities became evident in both neurites and somata of APP/PS1 mice at 4 months old, when senile plaques were only beginning to emerge [35] **(Fig. 3d)**. In contrast, activities of their coupled [Ca^2+^]_cyto_ were not significantly altered, suggesting that [Ca^2+^]_mito_ may be more sensitive to the pathological changes associated with AD (**Fig. 3e**).

**Figure 3.**
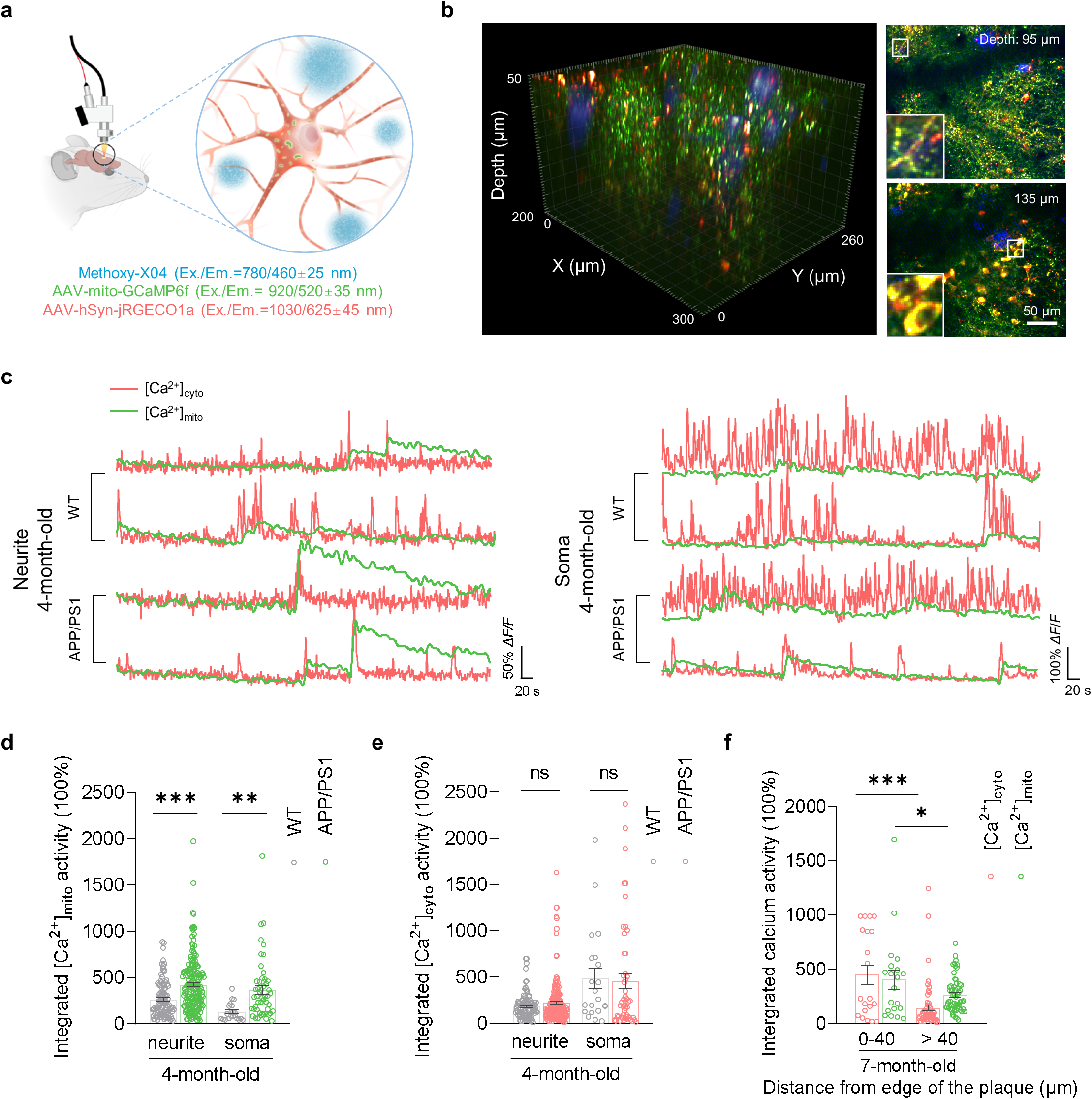
Multicolor imaging of [Ca^2+^]_mito_, [Ca^2+^]_cyto_ and amyloid plaques in APP/PS1 mice using FHIRM-TPM 3.0-U. **(a)** A schematic showing three-color imaging strategy. Amyloid plaques were labeled with methoxy-X04 and imaged at Ex./Em. = 780/460±25 nm. Mitochondrial Ca^2+^ was sparsely labeled with mito-DIO-GCaMP6f and CaMKII-Cre, imaged at Ex./Em. = 920/520±35 nm. Cytosolic Ca^2+^ was reported by hSyn-jRGECO1a and imaged at Ex./Em. = 1030/625±45 nm. Ex., excitation; Em., emission. **(b)** Multicolor imaging of amyloid plaques, mitochondria and neurons. Left, representative three-dimensional reconstruction from a 7-month-old APP/PS1 mouse. Right, three-color images at 95 μm (L1) and 135 μm (L2) depths below the pial surface. **(c)** Representative examples of pairs of [Ca^2+^]_cyto_ and [Ca^2+^]_mito_ transients in L1 neurites and L2 somas in 4-month-old WT and APP/PS1 mice. **(d, e)** Integrated activities of coupled [Ca^2+^]_mito_ (**d**) and [Ca^2+^]_cyto_ (**e**) in neurites and somas of 4-month-old WT and APP/PS1 mice. Data from 108 neurite events and 21 somatic events from 4 WT mice; 176 neurite events and 50 somatic events from 3 APP/PS1 mice. Integrated activities in d, WT neurites: 263.7±18.57; APP/PS1 neurites: 422.7±21.90; WT somas: 125.6±20.25; APP/PS1 somas: 366.1±47.10. Integrated activities in e, WT neurites: 182.8±12.83; APP/PS1 neurites: 219.1±18.17; WT somas: 486.9±111.6; APP/PS1 somas: 457.3±83.10. **(f)** Integrated activities of coupled [Ca^2+^]_cyto_ and [Ca^2+^]_mito_ were higher in the vicinity of a plaque (<=40 μm, 20 events, [Ca^2+^]_cyto_: 450.2±88.70; [Ca^2+^]_mito_: 405.8±88.07) compared to farther-away events (>40 μm, 59 events, [Ca^2+^]_cyto_: 142.1±27.94; [Ca^2+^]_mito_: 262.9±20.98) in APP/PS1 mice (n=3 at 7-month-old). Statistical analysis was performed using an unpaired t-test. Data are presented as mean ± SEM. *P < 0.05. **P < 0.01. ***P < 0.001.

Furthermore, to assess the association of this Ca^2+^ dysfunction with the deposition of amyloid plaques, the APP/PS1 mice were subsequently imaged at 7 months old. Interestingly, higher [Ca^2+^]_cyto_ and [Ca^2+^]_mito_ activities were observed proximal to the plaques (<=40 μm), indicating a local effect of amyloid plaques on both [Ca^2+^]_cyto_ and [Ca^2+^]_mito_ dynamics (**Fig. 3f**). Collectively, these results uncover an early dysregulation in [Ca^2+^]_mito_ dynamics and its potential link to amyloid plaque pathology in APP/PS1 mice.

## Discussion

FHIRM-TPM 3.0 presents a comprehensive solution to long-standing challenges of m2PMs. It enables multicolor achromatic imaging at 780 nm, 920 nm, and 1030 nm, and achieves functional and structural imaging at depths exceeding 820 μm. The interchangeable objective design allows for switching among different resolutions and FOVs, analogous to benchtop two-photon microscopy. We anticipate that this new system will become a versatile and powerful tool for exploring diverse research scenarios in neuroscience. With FHIRM-TPM 3.0 serving as a foundational platform, future developments in miniature multiphoton microscopy may emphasize strategies for higher speed, greater depth as well as the combination of imaging with precise optogenetic stimulation in freely behaving paradigms.

## Data availability

Two-photon imaging datasets acquired from animals are available upon request.

## Methods

### The FHIRM-TPM 3.0 system

The FHIRM-TPM 3.0 system was equipped with three femtosecond lasers with wavelengths of 780 nm (X-780, Carmel Laser, Inc., USA), 920 nm (AUTVS-AL15, SPARK LASERS, France) and 1030 nm (Seed-FL-1030-S1-80, Beijing Fatonics Technology Co., China). These lasers had a maximum average power of 1 to 1.5 W, a pulse width of approximately 100 fs, and a repetition rate of 80 MHz. An external half-wave plate (SAQWP05M-700, Thorlabs Inc., USA) and an Acousto-Optic Modulator (AOM) (MT110-B50A1.5-IR-Hk, AA Sa., Orsay Cedex, France) were placed in the respective optical paths to rapidly control the laser power. Tunable beam expanders (TVS-BEP-0.5/2×-B, Transcend Vivoscope Biotech Co., China) were used to adjust the beam diameter for efficient fiber coupling. The three laser beams were combined using a mirror and dichroic mirror set (DMSP820B and DMLP1000, Thorlabs Inc., USA), then coupled into an anti-resonant hollow-core fiber (AR-HCF) via a doublet (354171-B, LightPath Technologies, USA), and transmitted to the headpiece. The optical configuration of the headpiece was designed using Zemax software (OpticStudio 21.3.2, Zemax). A miniature collimator, consisting of a negative lens and a positive doublet, was mounted on the other end of AR-HCF, producing a collimated beam with a diameter of 1.2 mm at 780 nm transmission. The MEMS scanner (A3I12.2, Mirrorcle Technologies Inc., Richmond, CA, USA) featured a 1.2-mm diameter mirror, used at a 20° incidence angle, with the first resonant frequency of both axes at approximately 2.9 kHz. After reflecting off a mirror and the MEMS, the collimated beam passed through a scan lens with a large angle, a dichroic mirror, and was finally focused on the brain tissue by a miniature objective (U, HR, LF). By adjusting the 920 nm and 1030 nm laser dispersion, the positive dispersion induced by optical elements was compensated, achieving an exit dispersion close to 100 fs under the objective. The excited fluorescence was collected into a supple fiber bundle (SFB) by the miniature objective (U, HR, or LF) and an aspherical condenser, and then transmitted by the SFB to a photomultiplier tube (PMT) module. The effective optical diameter of the SFB was 1 mm, with a numerical aperture (NA) of 0.57. In the PMT module, the fluorescence was first collimated by a collimating lens and passed through a low-pass filter (SP-700-25, Transcend Vivoscope Biotech Co., China) to remove the stray laser light. Subsequently, blue, green, and red channels of fluorescence were separated by a dichroic mirror set (DM-SP496-25×36 and DM-SP556-25×36, Transcend Vivoscope Biotech Co., China), and directed into three PMTs through a band-pass filter set (BP-460/50-25, BP-520/70-25 and BP-625/90-25, Transcend Vivoscope Biotech Co., China), respectively.

### Animals

All the animal housing and experimental procedures were approved by the Peking University Animal Use and Care Committee and complied with the Association for Assessment and Accreditation of Laboratory Animal Care standards. Male C57BL/6 mice, Thy1-YFPH transgenic mice, APP/PS1 (APPswe/PSEN1ΔE9) transgenic mice (Jackson Laboratories, stock no. 034829) and age-matched wild-type littermate controls were used in these experiments. Animals were housed with a 12-hour light-dark cycle at 22 °C with 50% humidity and had free access to food and water.

### Virus injection

Mice were anesthetized with 1.5% isoflurane in the air at a flow rate of 0.4 L/min and mounted on a stereotactic frame. AAV viruses were slowly injected into targeted brain areas at a speed of 10 nL min^−1^ based on strategies listed in **Supplementary table 3**. The injection needle was kept in sites for another 5 minutes after injection.

### Cranial window preparation

For cortex imaging, one week after virus injection, a 4.3-mm-diameter craniotomy was made. Mice were anesthetized with isoflurane, and a piece of the skull (4.3 mm diameter) was carefully removed with the dura intact over the targeted cortex (**Supplementary table 3**). A piece of glass coverslip (4.3 mm diameter, TJ-001, 85– 115 μm, Optowide Technologies Inc.) was placed on the craniotomy and sealed with cyanoacrylate and dental cement.

### GRIN lens implant preparation

For hippocampus imaging, a GRIN lens was surgically implanted one week after virus injection. Mice were anesthetized and a 1.1 mm-diameter craniotomy was first made at the coordinates (AP: −2.2 mm, ML: −1.5 mm). After the skull was removed, the cortical tissue and callous corpus were aspirated using a 30-gauge blunted needle perpendicularly with the underlying dorsal CA1 intact. The needle was attached to a custom-constructed three-axis motorized robotic arm controlled by a MATLAB-based software [36]. When the entire area was clean, a 1-mm diameter GRIN lens (GoFoton) was slowly implanted above the dorsal hippocampal CA1 and secured to the skull using dental cement [37].

### Placement of the headpiece

The protocol for mounting, dismounting, and remounting the headpiece consisted of four steps, as described previously [9, 10]. Step 1: The baseplate with a coverslip was fixed over a cranial window with dental cement. Step 2: The headpiece was positioned. The headpiece was first screw-fastened to its holder and then placed onto the baseplate using a triaxial motorized stage. The triaxial motorized stage moved the headpiece precisely to focus on the desired depth. Once the region of interest was identified, the holder was cemented to the baseplate, allowing the mouse to move freely. Step 3: The headpiece was dismounted by being unscrewed and unplugged from the holder. Step 4: For remounting, a drop of water was applied to the coverslip of the baseplate, followed by the headpiece being plugged in and securely screw-fastened to the holder. In most experiments, we could relocate the same focal plane used previously.

### Three-color imaging in awake head-fixed APP/PS1 mice

To label amyloid plaques, methoxy-X04 (5 mg/kg) was intraperitoneally injected 24 hours before the imaging session. The two-photon excitation at 780 nm, 920 nm, 1030 nm and emission collection at 460±25 nm, 520±35 nm, 625±45 nm were used for methoxy-X04, mito-GCaMP6f and jRGECO1a, respectively. Three-color z-stack imaging was performed using a motorized stage to incrementally adjust the focal plane depth in 1-μm steps. Time-lapse imaging was conducted to capture Ca^2+^ signals at specific focal planes, consisting of 3,000 sequential frames. The average laser power of imaging was under 100 mw and the frame rate was 5 Hz.

### Scalable FOV imaging

After imaging with one of the three interchangeable parfocal objectives, the headpiece was unscrewed and unplugged from the holder. A subsequent objective was then installed, with a drop of water placed on the coverslip of the baseplate, and the headpiece was plugged in and screw-fastened to the holder. In most experiments, we could relocate the same region of interest, with FOV and resolution depending on the specific objective in use.

### Electric foot shock test

Mice were habituated for 3 minutes on a shocking grid under m2PM imaging, followed by manually scrambled electrical foot shocks (2 s, 0.4 mA) (**Extended Data Fig. 6a, b** and **Supplementary Video 1**). They were then observed for approximately 3 minutes post-shock.

### Data processing and analysis

We used the open-source software ImageJ for averaging, cropping, registration, pseudocolor-coding, and three-dimensional projection of raw images. All cross-section profiles, statistical graphs, traces of Ca^2+^ transients, and other coordinate graphs were drawn with GraphPad Prism 8 (GraphPad Software). All three-dimensional reconstruction figures and movies were made using Imaris (Bitplane).

Ca^2+^ imaging data of freely moving mice were analyzed offline with custom-written programs in MATLAB (MathWorks). We used Image Stabilizer (ImageJ) to correct motion artifacts before processing timeseries images. We then used an annular ring subtraction method [38] to present Ca^2+^ signals as relative fluorescence changes (ΔF/F). To analyze coupled [Ca^2+^]_cyto_ and [Ca^2+^]_mito_, regions of interest (ROIs) in green channel for imaging [Ca^2+^]_mito_ were segmented by suite2p [39] and the somata or neurites containing the same ROIs in red channel for imaging [Ca^2+^]_cyto_ were manually selected on the basis of morphology. Ca^2+^ signals were presented as relative fluorescence changes (ΔF/F), corresponding to changes in mean fluorescence from specified ROIs. The threshold for determining Ca^2+^ transients was calculated as three times the standard deviation (SD) of baseline fluorescence. Regions exhibited coupled [Ca^2+^]_cyto_ and [Ca^2+^]_mito_ were further analyzed. The integrated Ca^2+^ activity was the accumulated Ca^2+^ activities above the threshold. Plaque distance was the 3D Euclidean distance between the centroid of the mitochondrion ROIs and the nearest methoxy positive voxel.

## Acknowledgements

We thank Z. Zhao and H. Wu at the Institute of Basic Medical Sciences for their support with the biological experiments; L. Chen, J. Wang, D. Zhang, and H. Zhang at Peking University for their valuable comments on the manuscript and data processing; C. Wang and L. Feng at Beihang University for data processing. J. Tian, Y. Huang, and S. Wang at Beijing Transcend Vivoscope Biotech Co. for manuscript proofreading and mechanical and software support; and T. Zhao, H. Zhou and H. Fang at the Nanjing Brain Observatory for their biological experiment support and data-processing services. The work was supported by grants from the STI2030-Major Projects (2022ZD0212100 and 2022ZD0211903 for R.W., and 2021ZD0202205 for H.C.), the National Natural Science Foundation of China (32201130 for R.W., and 32293210 and 32327802 for H.C.), the National Key R&D Program of China (2022YFF1501900 for H. C., and 2023YFC3402604 and 2023YFF1501100 for A.W.), the CAMS Innovation Fund for Medical Sciences (2019-I2M-5-054) for H.C., the Beijing Natural Science Foundation (Z240013) for R.W., and China Postdoctoral Science Foundation (2023T160022) for C. Z..

## Author contributions

H.C., A.W., and R.W. conceived the project and supervised the research; R.W. designed and built the optical system; C.Z. designed the miniature optical configuration; S.Q. designed and performed the biological experiments under the supervision of X.W.; Y.Z. performed the experiments and figure-making; L.Z., Q.F., and H.H. performed the biological experiments; F.X. led the biological experiment data analysis; A.W. oversaw the fiber optics and assisted with the mechanical assembly; Y.H. provided support on the system under the supervision of Y.Z.; D.W. manufactured the optical fibers under the supervision of F.Y.; H.C., R.W., C.Z., A.W., S.Q., and Y.Z. wrote the manuscript; all authors participated in discussion and data interpretation.

## Competing interests

Q.F. and Y.H. are employees of the company Transcend Vivoscope, which develops and sells microscopes.

---

**Extended Data Fig. 1.**
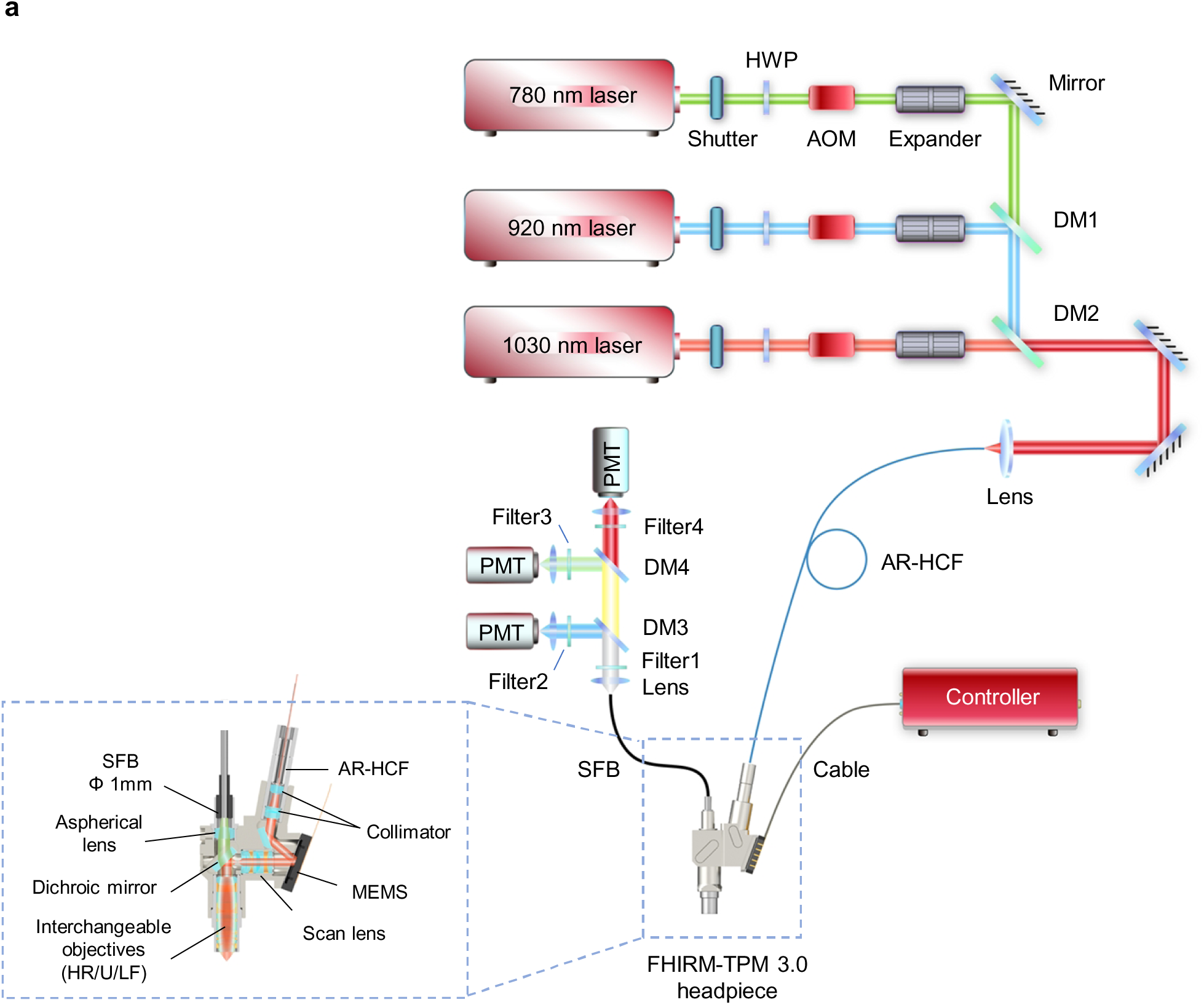
System design of FHIRM-TPM 3.0. **(a)** An overall scheme of FHIRM-TPM 3.0 with the inset displaying a profile of the headpiece. HWP, half-wave plate; AOM, acousto-optic modulator; DM, dichroic mirror; AR-HCF, anti-resonant hollow-core fiber; MEMS, microelectromechanical systems; SFB, supple fiber bundle; PMT, photomultiplier.

**Extended Data Fig. 2.**
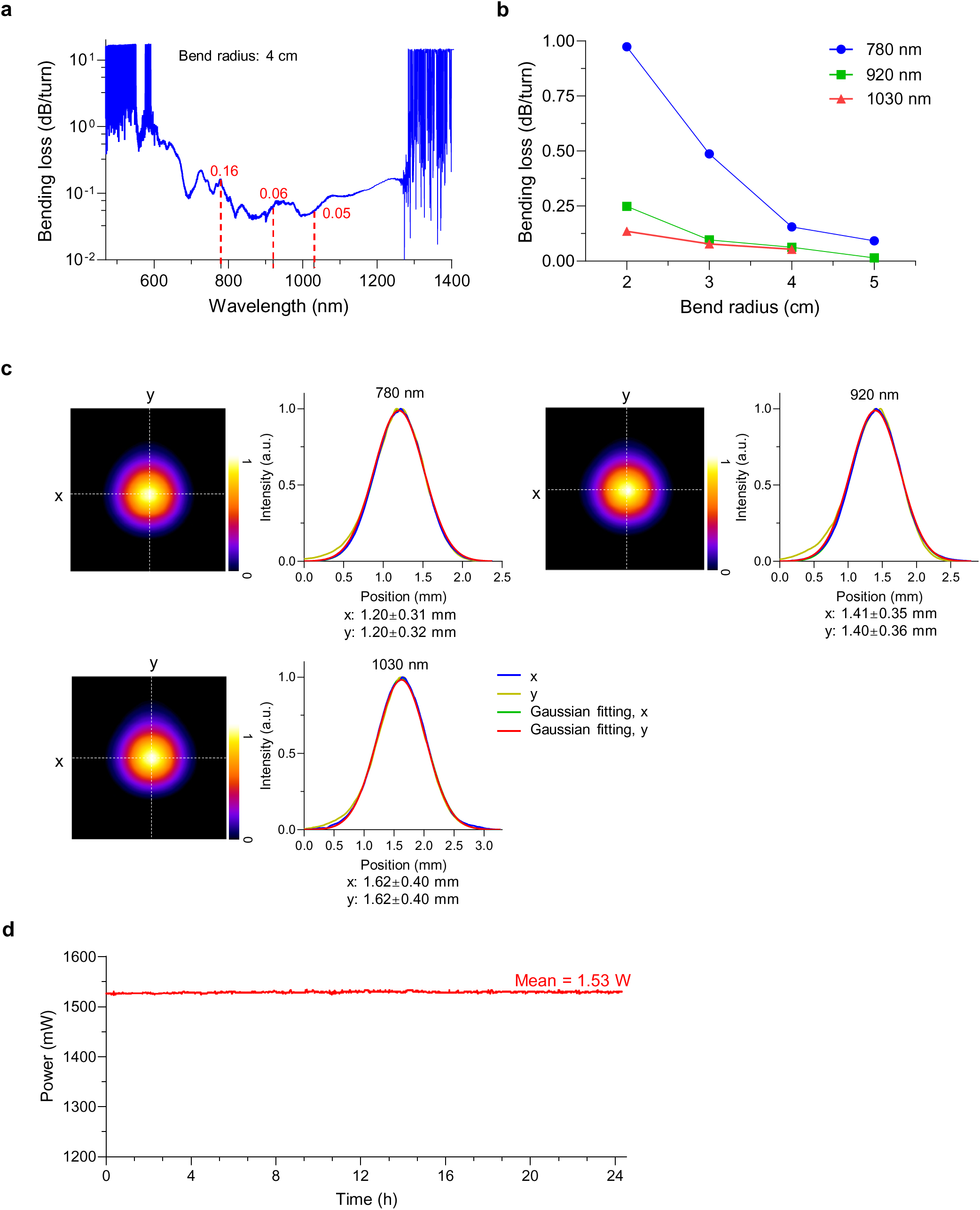
Performance test of the AR-HCF. **(a)** Bending loss as a function of wavelength for an AR-HCF with a bend radius of 4 cm. Red dashed lines indicate bending loss at 780 nm, 920 nm, and 1030 nm. **(b)** Bending loss as a function of bend radius in an AR-HCF at wavelengths of 780 nm, 920 nm, and 1030 nm, respectively. **(c)** Right, the x-y plots of the laser beam after AR-HCF at 780 nm, 920 nm and 1030 nm, respectively. Left, images of each laser beam delivered by AR-HCF after a collimator. Data are shown as mean ± SD. A.u., arbitrary units. **(d)** Stability test of high-power transmission. A 920 nm laser of 1.9 W (repetition rate: 80 MHz, pulse width: 149 fs) was coupled into a 2-m AR-HCF with an average output power of 1.53 W at the fiber exit. The output power was measured every second for 24.3 hours. The experiment was independently repeated n = 4 times.

**Extended Data Fig. 3.**
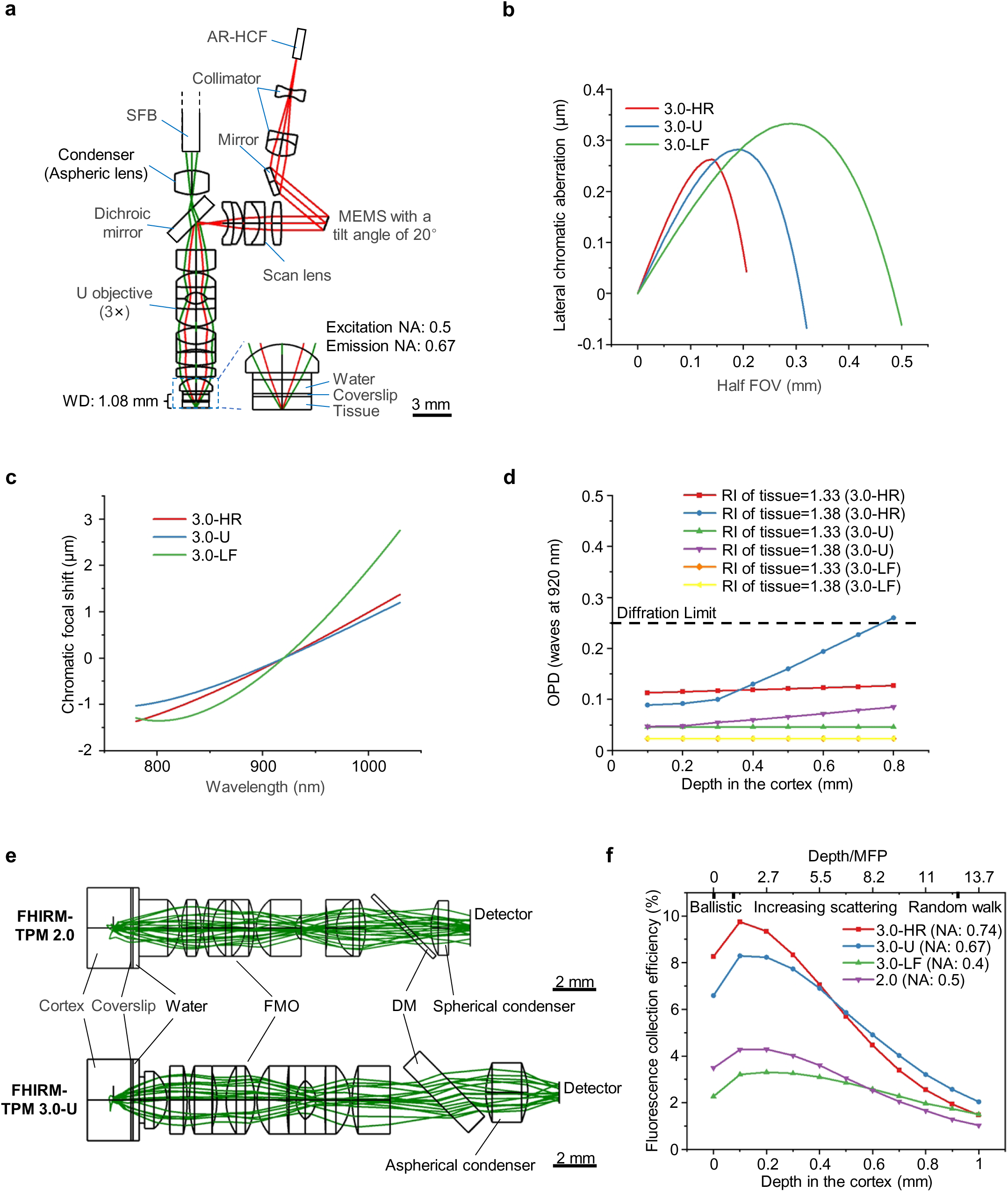
Optical design of the FHIRM-TPM 3.0 headpiece. **(a)** Optical layout of the FHIRM-TPM 3.0-U headpiece. AR-HCF, anti-resonant hollow-core fiber; SFB, supple fiber bundle; MEMS, microelectromechanical systems; WD, working distance. **(b)** Residual lateral chromatic aberrations in half the FOV across the waveband of 780 nm to 1030 nm for FHIRM-TPM 3.0-HR/U/LF. **(c)** Residual axial chromatic aberrations represented by chromatic focal shift across the waveband of 780 nm to 1030 nm. **(d)** Optical path difference (OPD) varying with imaging depth in simulated brain tissue with refractive indexes (RI) of 1.33 or 1.38. **(e)** Ray tracing diagram of scattered fluorescence for FHIRM-TPM 2.0 [10] and FHIRM-TPM 3.0-U. Green rays illustrate photons scattered in the cortical brain tissue, collected by the headpiece, and being directed onto the detector. FMO, finite miniature objective; DM, dichroic mirror. **(f)** Fluorescence collection efficiency as a function of imaging depth in simulated brain tissue for FHIRM-TPM 3.0-HR/U/LF compared with FHIRM-TPM 2.0. Note that as depth increases, the fluorescence is categorized into ballistic, scattering, and random walk regimes.

**Extended Data Fig. 4.**
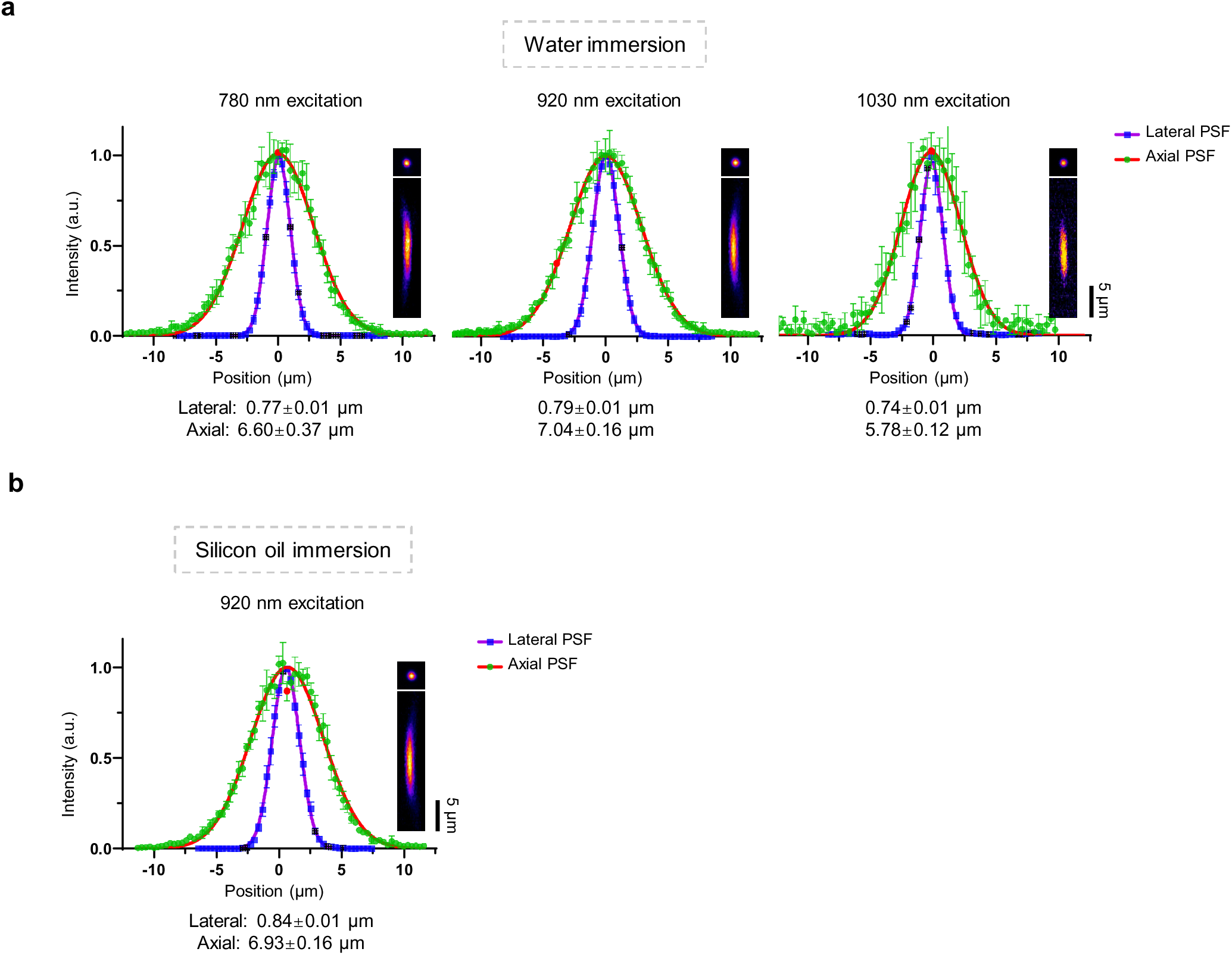
Resolution test of FHIRM-TPM 3.0-U (a, **b)** Resolution test of FHIRM-TPM 3.0-U. Normalized and averaged lateral (blue) and axial (green) profiles of four beads under 780 nm, 920 nm and 1030 nm excitation in water immersion **(a)**, as well as 920 nm excitation in oil immersion **(b)**. Insets, averaged x-y (top) and y-z (bottom) images of a 0.2 μm bead. PSF, point spread function. Data are shown as mean ± SEM.

**Extended Data Fig. 5.**
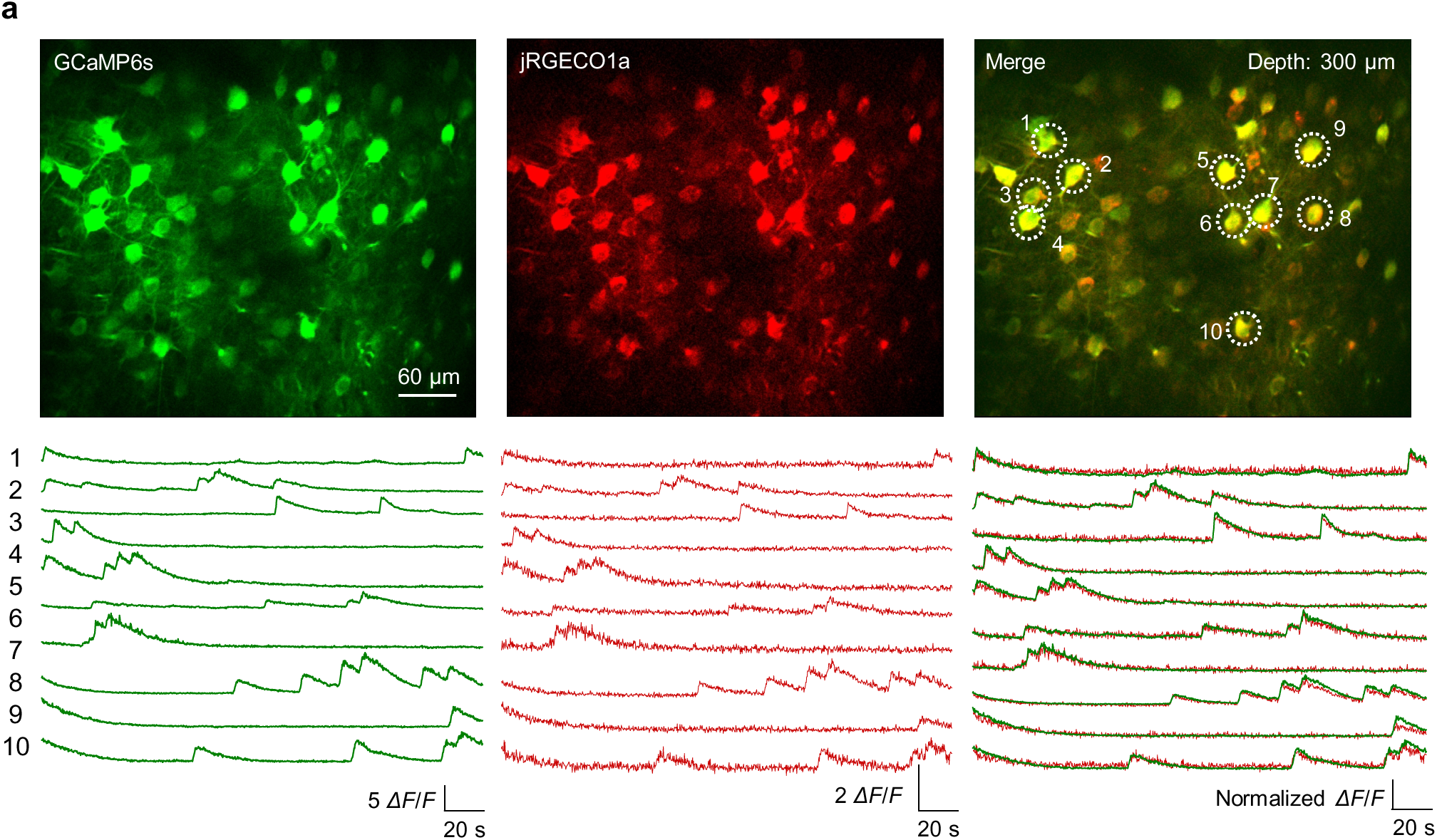
Dual-color imaging of neurons co-expressing GCaMP6s and jRGECO1a using FHIRM-TPM 3.0-U. **(a)** Top, average intensity projections of 200 frames from M1 neurons co-expressing GCaMP6s and jRGECO1a. Bottom, representative time courses of neuronal Ca²⁺ activity reported by GCaMP6s (green) and jRGECO1a (red), as marked by white circles in the top-right panel. Data were obtained from one animal and were representative of three similar experiments.

**Extended Data Fig. 6.**
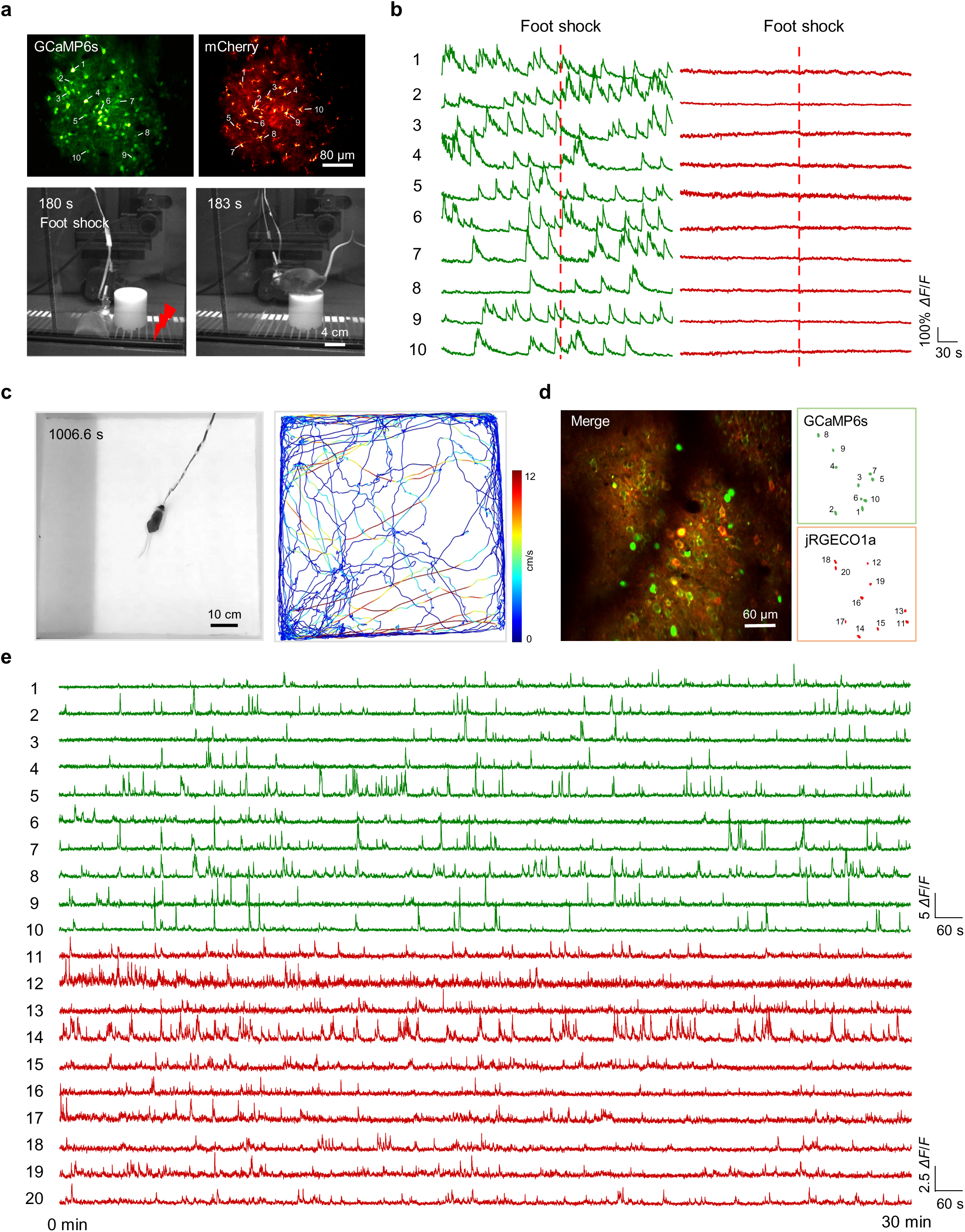
Dual-color imaging of mice experiencing electric foot shock and exploring an open field using FHIRM-TPM 3.0-U. **(a)** Stability test during dual-color imaging. Top images show neurons expressing GCaMP6s (left) and astrocytes expressing mCherry (right) in the M1 at depth of 300 μm, averaged over 100 frames. Bottom images show the mouse jumping from an electrical grid onto a platform during a 2-second foot shock. **(b)** Representative time courses of neuronal GCaMP6s-reported Ca^2+^ activity (green) and astrocytic mCherry fluorescence (red) in selected cells (marked with numerals in corresponding images). Dashed lines denote the foot shock and jumping. Data are representative of six experiments from six mice. **(c)** Left, representative snapshot of a mouse in an 80-cm square open-field box. Right, mouse trajectories extracted from a 30-minute test session video. Colors indicated movement speeds. **(d)** Left, average intensity projections of 100 frames from M1 neurons expressing GCaMP6s and jRGECO1a. Right, segmentation maps of twenty representative identified neurons. **(e)** Representative time courses of GCaMP6s (green) and jRGECO1a (red) reported neuronal Ca²⁺ activity in the segmented areas **(d)** (marked with numerals in corresponding images). Data were obtained from one animal and were representative of three similar experiments.

**Extended Data Fig. 7.**
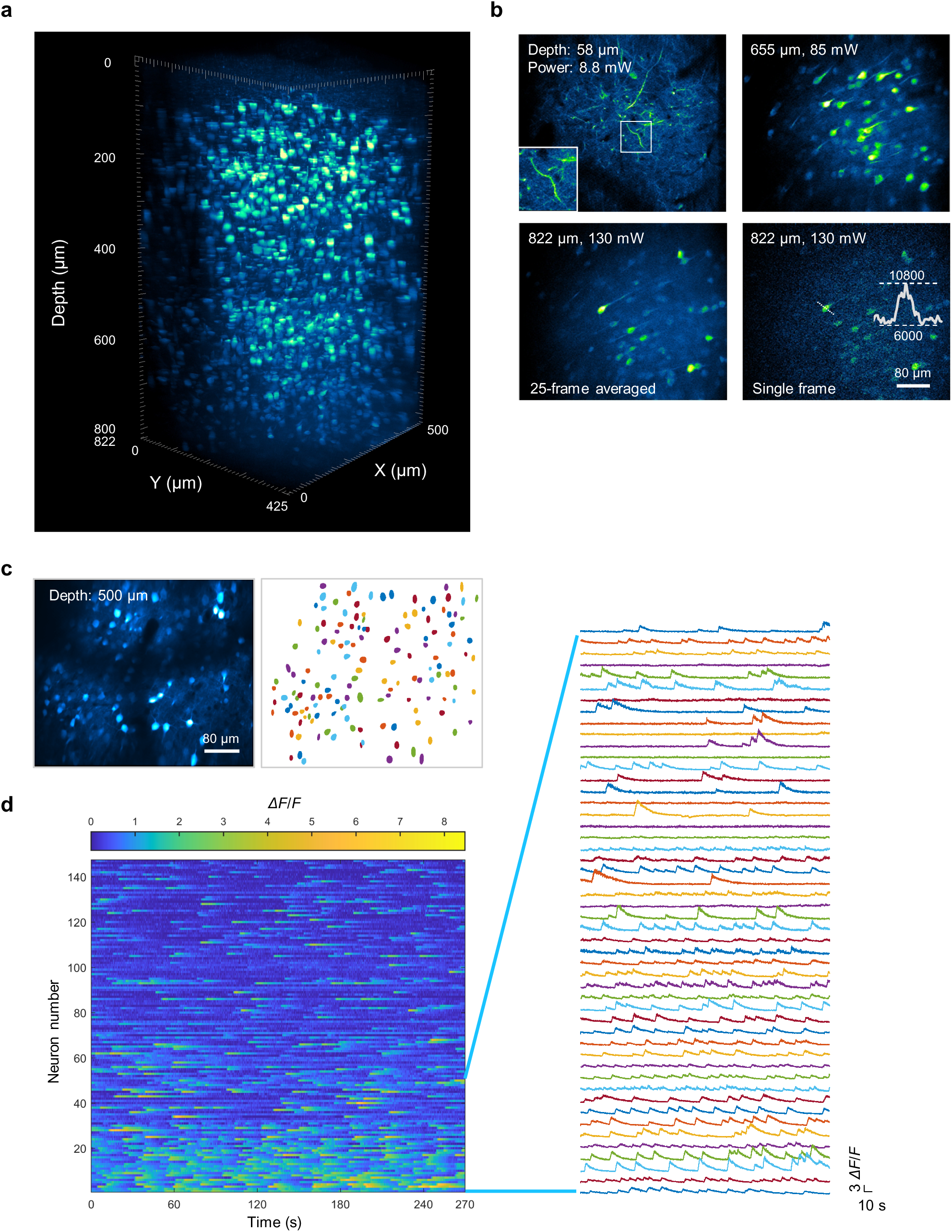
Deep-cortex Ca^2+^ imaging in freely moving mice using FHIRM-TPM 3.0-U. **(a)** Reconstruction of an 822 μm stack of GCaMP6s-labeled neurons in the mPFC of an awake mouse using 920 nm excitation with FHIRM-TPM 3.0-U. **(b)** Top and bottom left, representative 25-frame-averaged x–y images after motion correction at designated depths from the stack in **a**. Bottom right, a single-frame image at 822 μm depth with inset displaying the intensity profile along the cross line. The power values refer to the average laser power after the miniature objective. Data are representative of two experiments from two mice. **(c)** 100-frame-averaged image, after motion correction, showing neurons at a cortical depth of 500 μm (left) with segmentation map of 147 neurons (right) in a freely moving mouse. **(d)** Ca^2+^ activities of all 147 active neurons identified in **c**. Traces to the right show time course plots from 50 neurons. Data are representative of four experiments from four mice.

**Extended Data Fig. 8.**
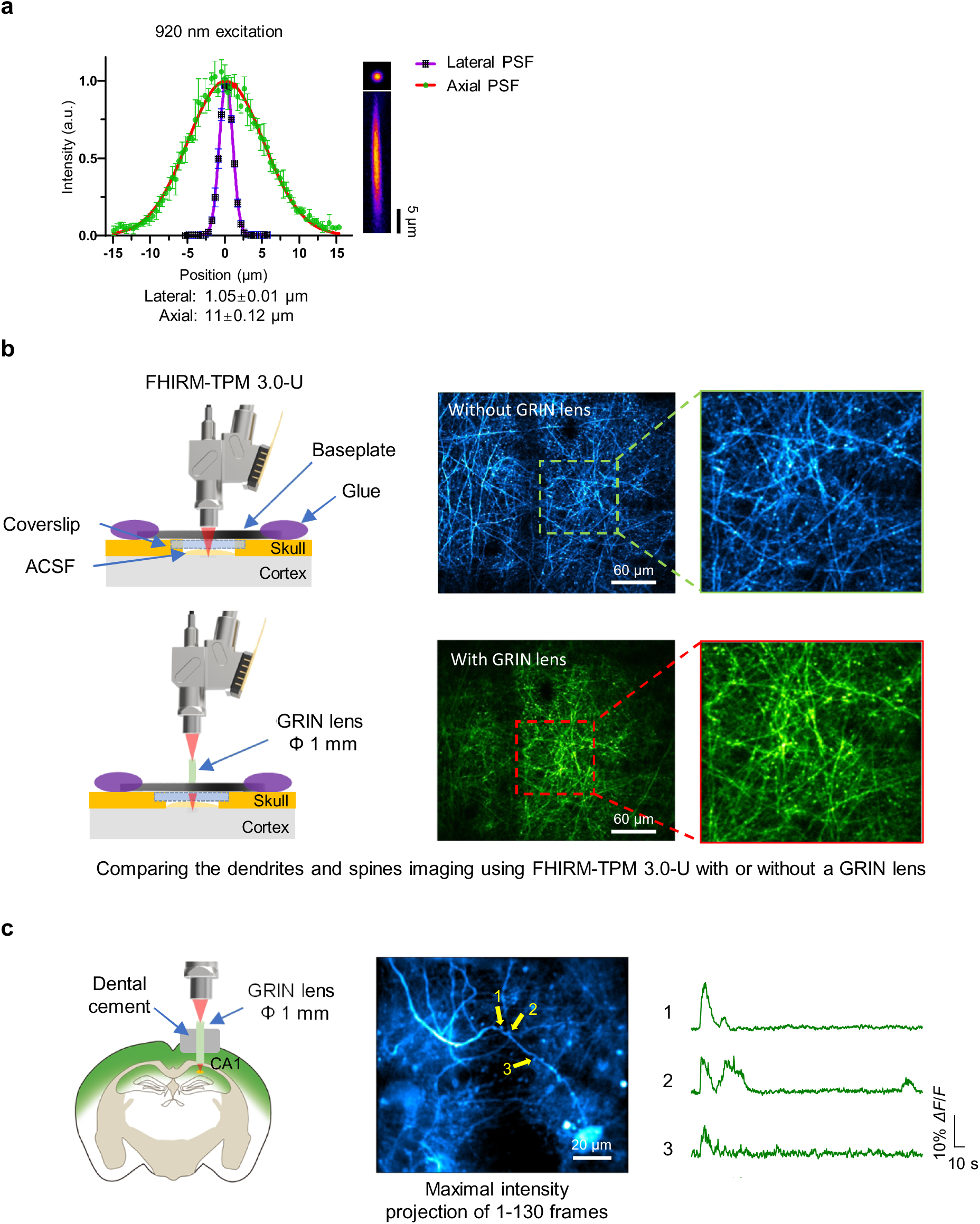
Using FHIRM-TPM 3.0-U with a GRIN lens to image hippocampal dendritic Ca^2+^ activities in vivo. **(a)** Resolution test of the FHIRM-TPM 3.0-U with a GRIN lens (1-mm diameter, 4-mm length). Left, normalized and averaged lateral (purple) and axial profiles (green) of four beads in water immersion under 920 nm excitation. Right, averaged x-y (top) and y-z (bottom) images of a 0.2 μm bead at 920 nm excitation. Data are shown as mean ± SEM. **(b)** Comparison of dendrites and spines imaging using the FHIRM-TPM 3.0-U with or without a GRIN lens. Left, a schematic of the two imaging methods. Right, superficial layer imaging of a Thy1-YFPH transgenic mouse in the M1 without (top, blue) or with a GRIN lens (1-mm diameter, 4-mm length, placed on the coverslip) (bottom, green). ACSF, artificial cerebrospinal fluid. The experiment was independently repeated n = 3 times. **(c)** Ca^2+^ imaging of hippocampus CA1 spines using the FHIRM-TPM 3.0-U through a GRIN lens (1-mm diameter, 4-mm length). Left, a schematic of the experimental setup. Middle, maximal intensity projection of 130 frames, after motion correction. Right, activities of GCaMP6s-labeled dendritic spines (arrows with numerals in the middle image). The experiment was independently repeated n = 2 times.

**Extended Data Fig. 9.**
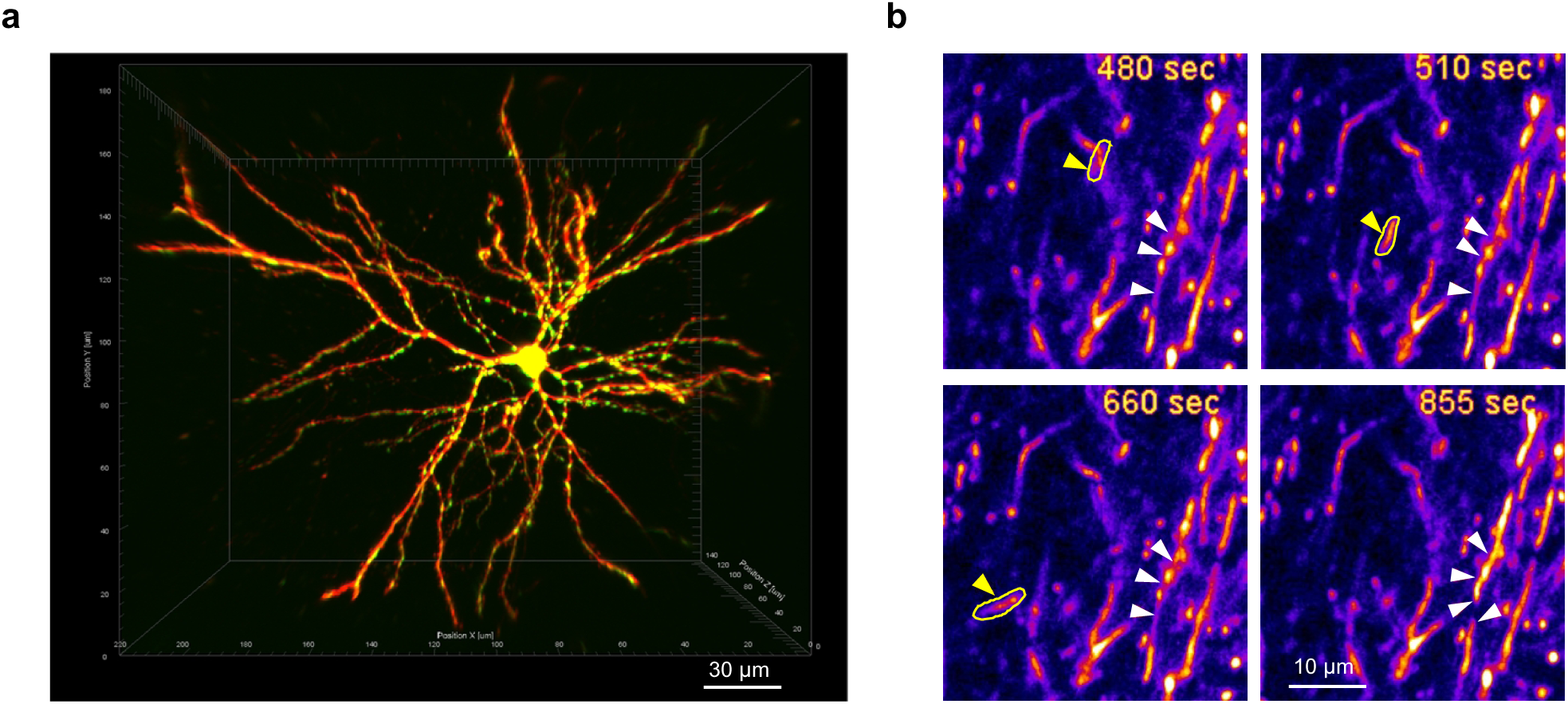
High-resolution imaging of mitochondrial motility and fission-fusion dynamics using FHIRM-TPM 3.0-HR. **(a)** Reconstruction of EGFP-labeled mitochondrial networks (920 nm excitation) in a tdTomato-labeled M1 neuron (1030 nm excitation) using FHIRM-TPM 3.0-HR. **(b)** Time-lapse images showing the movement of a mitochondrion (tracked by yellow arrows) and the fission-fusion dynamics of other mitochondria (tracked by array of white arrows).

**Extended Data Fig. 10.**
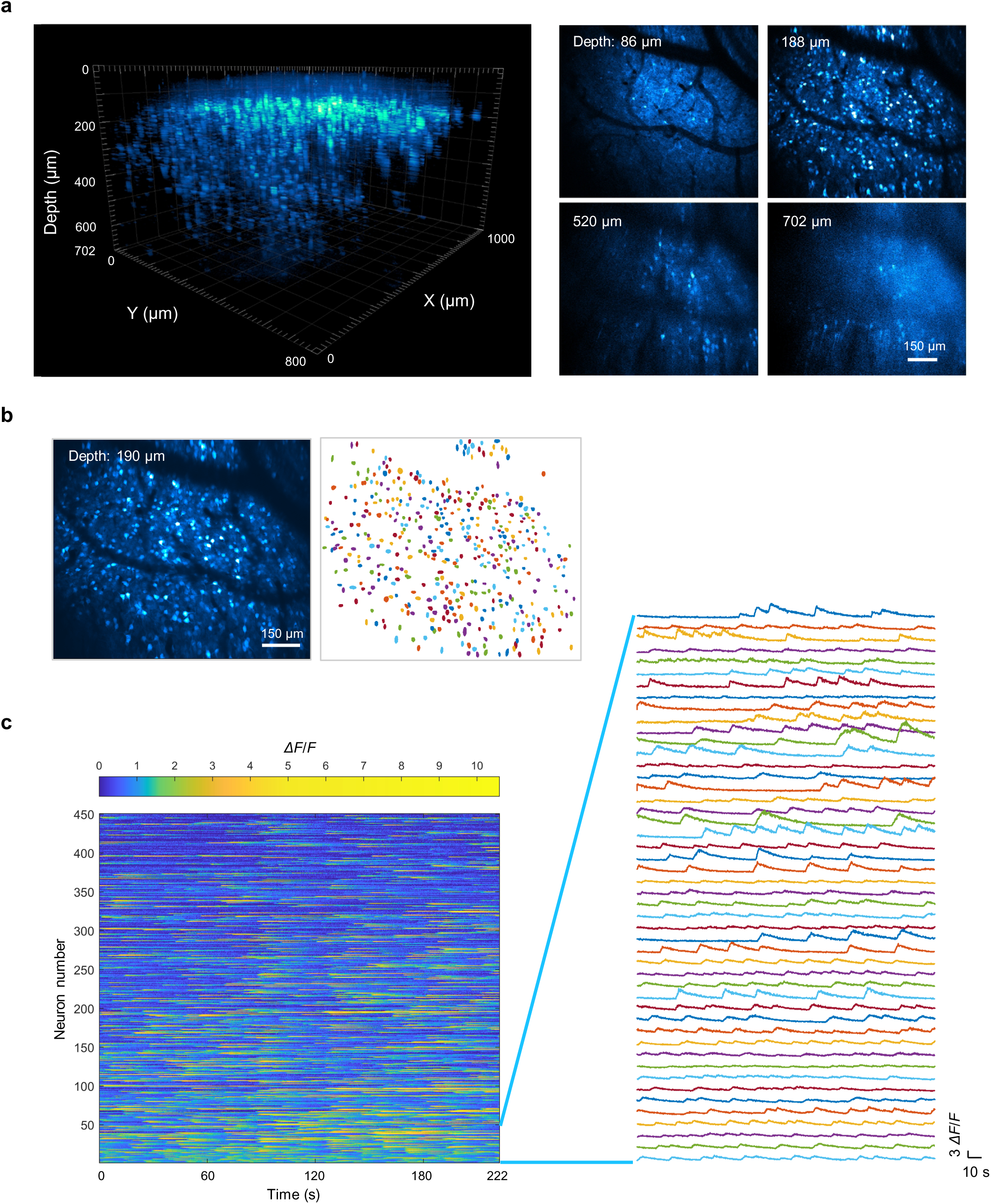
Millimeter FOV Ca^2+^ imaging with FHIRM-TPM 3.0-LF in a freely moving mouse. **(a)** Left, FHIRM-TPM 3.0-LF imaging of a stack (1.0×0.8×0.7 mm^3^) of GCaMP6s labeled neurons in the mPFC of an awake head-fixed mouse using 920 nm excitation. Right, representative 25-frame-averaged x–y images at designated depths from the left stack. The experiment was independently repeated n = 2 times. **(b)** A 100-frame-averaged image, after motion correction, at a depth of 190 μm (left) with its segmentation map of 450 neurons (right) from the mouse in free-moving conditions. **(c)** Ca^2+^ activities of all 450 neurons identified in **b**, with line plots for 50 neurons shown to the right. The experiment was independently repeated n = 4 times.

