## Supplementary Note for "A versatile miniature two-photon microscope enabling multicolor deep-brain imaging"

**Supplementary Note 1: Optical design**

The core optical design goals of FHIRM-TPM 3.0 are multicolor excitation, imaging depth extension, and switching between objectives of different magnification for scalable field-of-view (FOV). To achieve multi-color excitation, several sets of achromatic doublets were introduced in the collimator, scan lens and the headpiece objective (**Extended Data Fig. 3a**). By using various combinations of optical materials with different dispersions, the system achieved achromatism in the 780-1030 nm band. The residual lateral chromatic aberrations in half the FOV were less than 0.5 μm for FHIRM-TPM 3.0-HR/U/LF (**Extended Data Fig. 3b**). The residual axial chromatic aberrations for FHIRM-TPM 3.0-HR/U were approximately 1 μm in both the 780-920 nm and 920-1030 nm bands, while for FHIRM-TPM 3.0-LF, about 1 μm in the 780-920 nm band and 3 μm in the 920-1030 nm band (**Extended Data Fig. 3c**), the latter being only one-eighth of its axial PSF.

For deep brain imaging, several improvements were made in both fluorescence excitation and collection. Regarding fluorescence excitation, since the refractive index of the cortex (~1.38) is higher than that of water (~1.33), the spherical aberration increases with imaging depth in brain tissue. We therefore optimized the optical path of FHIRM-TPM 3.0-HR/U/LF, such that the spherical aberration introduced within the range of 0-800 μm cortical depth was below or slightly over the diffraction limit (**Extended Data Fig. 3d**). Regarding fluorescence collection, the optical path was separately designed to enlarge the collection numerical aperture (NA) for the HR, U, and LF objectives to 0.74, 0.67, and 0.4, respectively. Using the Henyey-Greenstein approximation to simulate brain tissue with a scattering length of 75 μm [40, 41] and an anisotropy parameter of 0.92 [41, 42], a ray-tracing model for the fluorescence collection path was established (**Extended Data Fig. 3e**). By adopting an aspheric condenser, the focus spot on the face of the collection supple fiber bundle (SFB) was reduced to 1 mm, allowing for finer fiber diameter to minimize obstruction to the free movement of mice. Monte Carlo tracing of 10 million rays showed that the fluorescence collection efficiency of FHIRM-TPM 3.0-U at an 800 μm imaging depth is twice that of FHIRM-TPM 2.0 (**Extended Data Fig. 3f**).

To enable imaging with scalable FOVs, we designed interchangeable parfocal objectives with different magnifications (**Fig. 1b**), HR: 4.65×, excitation NA: 0.7; U: 3×, excitation NA: 0.5; LF: 1.6×, excitation NA: 0.26. All three objectives have the same length, diameter, and working distance, allowing for interchangeability to meet the needs of balancing FOV and resolution, while facilitating the relocation of the same region and the same focal plane when switching between objectives.

Finally, we integrated the strategies recently developed to enlarge the FOV [26]. Specifically, the tilt angle of the scan MEMS was reduced to 20° to minimize compression of the scanning angle in the x direction. The scan lens with a large angle of ±15° was designed to match the maximum scanning angle of MEMS, attaining a millimeter FOV in FHIRM-TPM 3.0-LF.

### Supplementary Note 2: Evaluation of the GRIN lens impact on FHIRM-TPM 3.0-U imaging performance

To evaluate the effect of imaging through a GRIN lens, we acquired mouse cortical images at a depth of 30 μm in the same region, using FHIRM-TPM 3.0-U with and without a GRIN lens relay. We found that the resolution was moderately decreased by the use of a GRIN lens (lateral resolution: 0.79 μm versus 1.1 μm, axial resolution: 7.04 μm versus 11 μm) in addition to vignetting at the edges of the FOV (**Extended Data Fig. 8a, b**). Nonetheless, through a GRIN lens of 1-mm diameter and 4-mm length, we still achieved $Ca^{2+}$ imaging in dorsal hippocampal CA1 neurons at the single dendritic spine resolution (**Extended Data Fig. 8c**).

**Supplementary table 1: Specifications of AR-HCF**

| **AR-HCF** | |
|---|---|
| **Material** | |
| Fiber material | Pure silica |
| Coating material | Acrylate (single layer) |
| **Physical Properties** | |
| Core diameter | 28 μm |
| Cladding diameter | 200 μm |
| Coating diameter | 297 μm |
| **Optical Properties** | |
| Center wavelength | 780 nm |
| Transmission bandwidth (＜0.2 dB/m) | 500 - 1060 nm |
| Mode field diameter | 21.5 μm @780 nm<br>21.4 μm @ 920 nm<br>21.3 μm @1030 nm |
| Transmission loss | 0.08 dB/m @780 nm<br>0.10 dB/m @ 920 nm<br>0.14 dB/m @1030 nm |
| Estimated dispersion | 1.76 fs/nm/m @780 nm |

| | |
|---|---|
| | 2.19 fs/nm/m @920 nm<br>2.58 fs/nm/m @1030 nm |

**Supplementary table 2: Comparison of the FHIRM-TPM-3.0 with previous mTPMs**

| | Gibson Group 2018 [25] | Li Group 2021 [22] | MINI2P (D0213) 2022 [11]*3 | FHIRM-TPM 2017 [9] | FHIRM-TPM 2.0 2021 [10] | FHIRM-TPM 3.0-HR | FHIRM-TPM 3.0-U | FHIRM-TPM 3.0-LF |
|---|---|---|---|---|---|---|---|---|
| **Excitation wavelength (nm)** | 910 | 830/927(virtual)/1050 | 920 | 920 | 920 | 780/920/1030 | 780/920/1030 | 780/920/1030 |
| **Lateral resolution (μm)** | 2.6 | 0.8/0.9/1.0 | 1.15 | 0.65 | 1.13 | 0.63/0.68/0.71 | 0.77/0.79/0.74 | 1.35/1.46/1.59 |
| **Axial resolution (μm)** | 9.9 | 4.5/5.0/5.7 | 17.8 | 3.3 | 12.18 | 3.54/3.73/4.09 | 6.60/7.04/5.78 | 24.99/23.68/22.99 |
| **Imaging depth (μm)*1** | 180 | ~100 | 200 | 150 | 250 | 760 | 854 | 702 |
| **Working distance (μm)** | 350 | 200 | 1170 | 200 | 1000 | 1000 | 1000 | 1000 |
| **Maximum field of view ($\mu m^2$)** | Φ 240 | Φ 120 | ~500×500 /~420×420*2 | 130×130 | 420×420 | 300×260 | 500×425 | 1000×800 |
| **Maximum speed (Hz)** | 1.3-2.5 | 1.5 | 15/40 (256 lines) *2 | 40 (256 lines) | 20 (256 lines) | 20 (256 lines) | 20 (256 lines) | 20 (256 lines) |
| **Scanning mechanism** | Galvo with fiber buddle | PZT | MEMS | MEMS | MEMS | MEMS | MEMS | MEMS |
| **Weight (g)** | ~2.5 (3D) | ~1 | 2.4 (3D) | 2.15 | 2.45/ 4.2 (3D) | 2.6 | 2.6 | 2.6 |
| **Immersion medium** | Air | Water | Water | Water | Water | Water/Silicon oil | Water/Silicon oil | Water/Silicon oil |

| | | | | | | | | |
|---|---|---|---|---|---|---|---|---|
| **Free-moving experiment** | Mouse | Not shown | Mouse | Mouse | Mouse | Mouse | Mouse | Mouse |
| **Indicators** | GCaMP6<br>tdTomato | CFP<br>GFP<br>YFP<br>RFP | GCaMP<br>tdTomato | GCaMP6<br>GFP | GCaMP6 | methoxy-X04<br>GCaMP6<br>mitoEGFP<br>YFP<br>tdTomato<br>jRGECO<br>mCherry | methoxy-X04<br>GCaMP6<br>mitoEGFP<br>YFP<br>tdTomato<br>jRGECO<br>mCherry | methoxy-X04<br>GCaMP6<br>mitoEGFP<br>YFP<br>tdTomato<br>jRGECO<br>mCherry |

*1: Derived from the figures in the paper

*2: Results of different MEMS configurations

*3: D0213 indicates the model of the objective equipped on the MINI2P

**Supplementary table 3: Summary of transgenic mice, dyes and virus used for in vivo imaging**

| Figure | Aim | Transgenic/virus/dye | Titer and volume/ mouse | Brain area | Stereotactic coordinates |
|---|---|---|---|---|---|
| **Fig. 2a-c, Extended Data Fig. 8b** | Neuronal structure | Thy1-YFPH | / | M1 | / |
| **Fig. 2d, Fig. 3** | Amyloid plaques | methoxy-X04 | 5 mg/kg | M1 | AP: 0.5 mm<br>ML: 1.2 mm<br>DV: -0.3 mm |
| | Mitochondrial calcium signals | AAV-EF1AcDIO-mito-GCaMP6f-WPRE | $5\times10^{12}$ vg/mL, 100 nL | | |
| | | rAAV-CaMKII-Cre-WPRE-pA (diluted 100-fold with PBS) | $5\times10^{12}$ vg/mL, 50 nL | | |
| | Neuronal calcium signals | rAAV-hSyn-NES-jRGECO1a-WPRE-hGH-pA | $5\times10^{12}$ vg/mL, 200 nL | | |
| **Extended Data Fig. 5** | Neuronal calcium signals | AAV-hSyn-GCaMP6s | $2.5\times10^{12}$ vg/mL, 150 nL | | |
| | | AAV-hSyn-jRGECO1a | $2.5\times10^{12}$ vg/mL, 150 nL | | |
| **Extended Data** | Neuronal | AAV-hSyn-GCaMP6s- | $2.5\times10^{12}$ vg/mL, | | |

| | | | | | |
|---|---|---|---|---|---|
| **Fig. 6a** | calcium signals | WPRE-hGH-pA | 200 nL | | |
| | Astrocytes structure | rAAV-GfaABCID-mCherry-WPRE-SV40pA | $2.5\times10^{12}$ vg/mL, 150 nL | | |
| **Extended Data Fig. 6d** | Neuronal calcium signals | AAV-hSyn-GCaMP6s | $2.5\times10^{12}$ vg/mL, 150 nL | | |
| | | AAV-CaMKII-jRGECO1a | $2.5\times10^{12}$ vg/mL, 150 nL | | |
| **Extended Data Fig. 9a, b** | Neuronal and mitochondrial structure | AAV-hSyn-DIO-mitoEGFP-2A-tdTomato | $5\times10^{12}$ vg/mL, 150 nL | | |
| | | rAAV-CaMKII-Cre-WPRE-pA (diluted 500-fold with PBS) | $5\times10^{12}$ vg/mL, 150 nL | | |
| **Fig. 2e, Extended Data Fig. 7a-c, Extended Data Fig. 10a, b** | Neuronal calcium signal | AAV-hSyn-GCaMP6s-WPRE-hGH-pA | $5\times10^{12}$ vg/mL, 500 nL | mPFC | AP: 2.8 mm<br>ML: 0.33 mm<br>DV: -0.4 mm |
| **Extended Data Fig. 8c** | | | $3\times10^{11}$ vg/mL, 200 nL | CA1 | AP: -2.2 mm<br>ML: -1.5 mm<br>DV: - 1.3 mm |

### Supplementary table 4: Summary of imaging parameters

| | **Fig. 2a-c Supplementary Video 2** | **Fig. 2d** | **Fig. 2e** | **Fig. 3b, Supplementary Video 10** | **Extended Data Fig. 5** |
|---|---|---|---|---|---|
| **Experimental paradigm** | Head-fixed | | | Head-fixed | Head-fixed |
| **Indicators** | Thy1-YFPH | GCaMP6f | GCaMP6s | methoxy-X04<br>mito-GCaMP6f<br>jRGECO1a | GCaMP6s<br>jRGECO1a |
| **Imaging volume (xyzt) (µm×µm×µm ×s)** | 500×425×854×373.8<br>(7 frames/plane) | 717×616×1×200<br>410×352×1×200<br>253×218×1×200 | 300×260×1×222.2<br>500×425×1×222.2<br>1000×800×1×222.2 | 296×254×166×1328<br>(40 frames/plane) | 450×390×1×200 |
| **Frame size (xyzt)** | 1200×1024×534×3738 | 512×440×1×1000 | 600×512×1×1000<br>600×450×1×1000 | 460×410×166×6640 | 590×512×1000 |

| | | | | | |
|---|---|---|---|---|---|
| | | | 512×420×1×1000 | | |
| **Frame rate (Hz)** | 10 | 5 | 4.5 | 5 | 5 |
| **Laser wavelength (nm)** | 920 for **Fig. 2a, b**<br>1030 for **Fig. 2c** | 920 | 920 | 780<br>920<br>1030 | 920<br>1030 |
| **Laser power (mW)** | 25-110 for **Fig. 2a**<br>66-110 for **Fig. 2b**<br>59-67 for **Fig. 2c** | 88 | 30 | 25 at 780 nm<br>30 at 920 nm<br>45 at 1030 nm | 30 at 920 nm<br>40 at 1030 nm |
| **Emission filter (nm)** | 520±35 | 520±35 | 520±35 | 460±25<br>520±35<br>625±45 | 520±35<br>625±45 |

| | **Extended Data Fig. 6a, Supplementary Video 1** | **Extended Data Fig. 6d** | **Extended Data Fig. 7a, b, Supplementary Video 3** | **Extended Data Fig. 7c, Supplementary Video 4** | **Extended Data Fig. 8b** |
|---|---|---|---|---|---|
| **Experimental paradigm** | Electric foot shock | Open-filed exploration | Head-fixed | Free-moving | Head-fixed |
| **Indicators** | GCaMP6s<br>mCherry | GCaMP6s<br>jRGECO1a | GCaMP6s | | Thy1-YFPH |
| **Imaging volume (xyzt) (μm×μm×μm×s)** | 500×425×1×350 | 420×420×1×1800 | 500×425×822×1370<br>(25 frames/plane) | 500×425×1×270 | 320×280×1×0.5 |
| **Frame size (xyzt)** | 600×512×1×3000 | 512×512×9000 | 600×512×548×13700 | 600×512×1×2700 | 940×830×1×5 |
| **Frame rate (Hz)** | 8.6 | 5 | 10 | 10 | 10 |
| **Laser wavelength (nm)** | 920<br>1030 | 920<br>1030 | 920 | 920 | 920 |
| **Laser power (mW)** | 35 at 920 nm<br>45 at 1030 nm | 35 at 920 nm<br>50 at 1030 nm | 8.8-130 | 65 | 12 |
| **Emission filter (nm)** | 520±35<br>625±45 | 520±35<br>625±45 | 520±35 | 520±35 | 520±35 |

| | **Extended Data Fig. 8c, Supplementary Video 5** | **Extended Data Fig. 9a, Supplementary Video 6** | **Extended Data Fig. 9b,** | **Extended Data Fig. 10a, Supplementary Video 8** | **Extended Data Fig. 10b, Suppleme** |
|---|---|---|---|---|---|

| | | | | | |
|---|---|---|---|---|---|
| | | | **Supplement ary Video 7** | | **ntary Video 9** |
| **Experimental paradigm** | Head-fixed | | | | Free-moving |
| **Indicators** | GCaMP6s | mito-EGFP tdTomato | mito-EGFP | GCaMP6s | |
| **Imaging volume (xyzt) (μm×μm×μm×s)** | 148×120×1×1 15.1 | 220×188×148×1 480 (100 frames/plane) | 42×44×1×15 29.9 | 1000×800×70 2×1775 (25 frames/plane) | 1000×800 ×1×222.2 |
| **Frame size (xyzt)** | 559×457×1×1 070 | 600×512×74×74 00 | 325×337×1× 7328 | 600×512×352 ×8775 | 600×512× 1×1000 |
| **Frame rate (Hz)** | 9.3 | 5 | 4.79 | 5 | 4.5 |
| **Laser wavelength (nm)** | 920 | 920<br>1030 | 920 | 920 | 920 |
| **Laser power (mW)** | 30 | 60 at 920nm<br>65 at 1030nm | 68 | 15-130 | 30 |
| **Emission filter (nm)** | 520±35 | 520±35<br>625±45 | 520±35 | 520±35 | 520±35 |


### Supplementary Video 1

Dual-color imaging of GCaMP6s-labeled neuronal $Ca^{2+}$ activity (920 nm excitation) and mCherry-labeled astrocyte morphology (1030 nm excitation) in the M1 during an electric foot shock experiment using the FHIRM-TPM 3.0-U at a frame rate of 8.3 Hz. On the right, representative traces of 10 selected neurons and 10 selected astrocytes are shown, with the red dashed line indicating the time points of electrical shocks. The moving white line on the traces is synchronized with the behavior video and neuronal activities. The video is shown at a fast speed (×2.8).

**Supplementary Video 2**

3D imaging in the M1 of an awake, head-fixed Thy1-YFPH transgenic mouse using the FHIRM-TPM 3.0-U. The imaged volume was 500×425×854 $\mu m^3$. 7 frames were averaged for each layer with an interval of 1.6 μm.

**Supplementary Video 3**

3D imaging of GCaMP6s-labeled neurons in the mPFC of an awake, head-fixed mouse using the FHIRM-TPM 3.0-U. The imaged volume was 500×425×822 $\mu m^3$. 25 frames were averaged after motion correction for each layer with an interval of 1.5 μm.

**Supplementary Video 4**

Time-lapse imaging of GCaMP6s-labeled neurons in a freely moving mouse using the FHIRM-TPM 3.0-U at a frame rate of 10 Hz. Data were obtained at 500 μm depth below the pial surface. The data were corrected for motion artifacts using the Image Stabilizer plugin (ImageJ) before finally being projected after 4-frame running average.

The video is shown at a fast speed (×3).

**Supplementary Video 5**

Time-lapse imaging of GCaMP6s-labeled hippocampus CA1 dendrite spines using the FHIRM-TPM 3.0-U through a GRIN lens (1-mm diameter, 4-mm length) at a frame rate of 9.3 Hz. The data were corrected for motion artifacts using the Image Stabilizer plugin (ImageJ) before finally being projected after 4-frame running average. The video is shown at a fast speed (×3).

**Supplementary Video 6**

3D imaging of sparsely labeled mitochondria expressing EGFP (920 nm excitation) in a neuron expressing tdTomato (1030 nm excitation) in the M1 of an awake, head-fixed mouse using the FHIRM-TPM 3.0-HR. The imaged volume was 220×188×148 $\mu m^3$. 100 frames were averaged for each layer with an interval of 2 μm.

**Supplementary Video 7**

Time-lapse imaging of EGFP-labeled mitochondria showing mitochondrial motion and fusion and fission dynamics using the FHIRM-TPM 3.0-HR. The yellow arrows indicated mitochondrial motion and purple arrows indicated mitochondrial fission and fusion dynamics. Data were obtained at 50 μm depth below the pial surface. The data were corrected for motion artifacts before being projected with the Image Stabilizer plugin (ImageJ).

**Supplementary Video 8**

3D imaging of GCaMP6s-labeled neurons in the mPFC of an awake, head-fixed mouse using the FHIRM-TPM 3.0-LF. The imaged volume was 1.0×0.8×0.7 $mm^3$. 25 frames were averaged for each layer with an interval of 2 μm.

**Supplementary Video 9**

Time-lapse imaging of GCaMP6s-labeled neurons in a freely moving mouse using the FHIRM-TPM 3.0-LF at a frame rate of 4.5 Hz. Data were obtained at 190 μm depth below the pial surface. The data were corrected for motion artifacts using the Image Stabilizer plugin (ImageJ) before finally being projected after 4-frame running average. The video is shown at a fast speed (×6.7).

**Supplementary Video 10**

3D imaging of methoxy-X04-labeled amyloid plaques (780 nm excitation), GCaMP6f-labeled mitochondria (920 nm excitation) and jRGECO1a-labeled neurons (1030 nm excitation) in the M1 of an awake, head-fixed mouse using the FHIRM-TPM 3.0-U. The imaged volume was 296×254×166 $\mu m^3$. 40 frames were averaged for each layer with an interval of 1 μm.
